# Unexpected organellar locations of ESCRT machinery in Giardia intestinalis and complex evolutionary dynamics spanning the transition to parasitism in the lineage Fornicata

**DOI:** 10.1101/2021.02.11.430673

**Authors:** Shweta V. Pipaliya, Rui Santos, Dayana Salas-Leiva, Erina A. Balmer, Corina D. Wirdnam, Andrew J. Roger, Adrian B. Hehl, Carmen Faso, Joel B. Dacks

**Affiliations:** Division of Infectious Diseases, Department of Medicine, University of Alberta, Edmonton, Alberta, Canada; Institute of Parasitology, University of Zurich, Zurich, Switzerland; Centre for Comparative Genomics and Evolutionary Bioinformatics, Department of Biochemistry and Molecular Biology, Faculty of Medicine, Dalhousie University, Halifax, Nova Scotia, Canada; Institute of Cell Biology, University of Bern, Bern, Switzerland; Institute of Parasitology, Biology Centre, CAS, v.v.i. Branisovska 31 370 05 Ceske Budejovice, Czech Republic

**Keywords:** ESCRT/Endosomal Sorting Complexes Required for Transport, VPS/Vacuolar Protein Sorting, PV/peripheral vacuoles, MVB/Multivesicular bodies, CHMP/Charged Multivesicular Protein, Fornicata, *Giardia*

## Abstract

Comparing a parasitic lineage to its free-living relatives is a powerful way to understand how the evolutionary transition to parasitism occurred. *Giardia intestinalis* (Fornicata) is a leading cause of gastrointestinal disease world-wide and is famous for its unusual complement of cellular compartments, such as having peripheral vacuoles instead of typical endosomal compartments. Endocytosis plays an important role in *Giardia*’s pathogenesis. Endosomal sorting complexes required for transport (ESCRT) are membrane-deforming proteins associated with the late endosome/multivesicular body (MVB). MVBs are ill-defined in *G. intestinalis* and roles for identified ESCRT-related proteins are not fully understood in the context of its unique endocytic system. Furthermore, components thought to be required for full ESCRT functionality have not yet been documented in this species.

We used genomic and transcriptomic data from several Fornicata species to clarify the evolutionary genome streamlining observed in *Giardia*, as well as to detect any divergent orthologs of the Fornicata ESCRT subunits. We observed differences in the ESCRT machinery complement between *Giardia* strains. Microscopy-based investigations of key components of ESCRT machinery such as *Gi*VPS36and *Gi*VPS25 link them to peripheral vacuoles, highlighting these organelles as simplified MVB equivalents. Unexpectedly, we show ESCRT components associated with the Endoplasmic Reticulum, and for the first time, mitosomes. Finally, we identified the rare ESCRT component CHMP7 in several fornicate representatives, including *Giardia*, and show that contrary to current understanding, CHMP7 evolved from a gene fusion of VPS25 and SNF7 domains, prior to the last eukaryotic common ancestor, over 1.5 billion years ago. Our findings show that ESCRT machinery in *G. intestinalis* is far more varied and complete than previously thought, and associating to multiple cellular locations and presenting changes in ESCRT complement which pre-date adoption of a parasitic lifestyle.

## INTRODUCTION

The food and waterborne diarrheal disease known as Giardiasis causes global healthcare and agricultural burden with approximately 300 million and more than 10 million cases diagnosed in humans and animals every year, respectively (Lanata et al., 2013). The causative agent is the diplomonad *Giardia intestinalis.* This enteric protist parasite has undergone large genome streamlining and modifications in its typical eukaryotic organelles, particularly in its endomembrane system and the associated trafficking complement (Faso and Hehl, 2011).

*Giardia* relies heavily on its endomembrane trafficking system to secrete virulence factors while establishing gut infection (Allain and Buret, 2020, Faso et. al., 2019), performing antigenic variation for immune system evasion (Gargantini et. al., 2016) and interfering with immune responses by degrading or reducing synthesis of signalling molecules (Ekmann et. al., 2000, Stadelmann et. al., 2012). Endomembrane trafficking is also required for completion of the life cycle during encystation which features regulated secretion of large amounts of cyst wall material through COPII- and COPI- associated lineage specific encystation specific vesicles (ESVs) (Stefanic et. al., 2009). *Giardia*’s endomembrane organization is significantly reduced in its complexity, most notably, because it lacks a canonical Golgi apparatus, readily identifiable early and late endosomes, lysosomes, and peroxisomes (Sheffield and Bjorvatn, 1977, Abodeely et al., 2009). Simplification of the endocytic and secretory pathways in this organism is underlined by complete loss of several protein complexes associated with membrane trafficking such as AP3, AP4, AP5, TSET, and the protein complexes that are present are often reduced in their complement such as Rabs, Rab GEFs, SNAREs, and ARF GEFs (Elias et al., 2012, Hirst et al., 2014, Venkatesh et al., 2017, Herman et al., 2018, Pipaliya et al., 2019). However*, Giardia* does harbour a tubulovesicular Endoplasmic reticulum (ER) thought to carry out functions of the late endosomal pathway (Abodeely et al., 2009). *Giardia* also has endocytic organelles called peripheral vacuoles (PVs) which perform bulk flow uptake of nutrients from the host environment and cargo sorting for retrograde transport (Zumthor et al., 2016; Cernikova et. al., 2020).

Endosomal Sorting Complexes Required for Transport (ESCRTs) are evolutionarily ancient complexes composed of five sub-complexes, ESCRT-0, ESCRTI, II, III, and III-A and recruited onto the growing late endosomal surface in a sequential manner to induce intraluminal vesicle formation through negative membrane deformation (Raiborg and Stenmark, 2009; Supplementary figure S1). In model eukaryotes, ESCRT machinery is required for the biogenesis of multivesicular bodies (MVBs) which have endocytic characteristics and the ability to mediate exosome biogenesis and release (Vietri et al., 2019). Nonetheless, additional ESCRTs functions are being discovered in plasma membrane repair, autophagy functions, post-mitotic nuclear envelope scission, and others with a shared function in membrane abscission (Hurley et al., 2015, Vietri et al., 2019). This conserved protein complex is never completely lost by organisms, underlining its importance, and was already elaborated in the LECA, presumably inherited from the Asgard archaea (Leung et. al., 2008, Spang et al., 2015, Seitz et al., 2019). Previous bioinformatics studies have shown *Giardia intestinalis assemblage* AI isolate AWB to possess patchy ESCRT-II, ESCRT-III, and ESCRT-IIIA machinery (Leung et al., 2008). However, key components within each of these were reported to be absent (Leung et al., 2008, Dutta et. al., 2015, Saha et al., 2018).

A powerful approach to understanding the evolutionary path to parasitism is to compare protein complements in parasites with those of free-living relatives. *Carpediemonas membranifera* is a small heterotrophic flagellate, and the namesake for the paraphyletic group of free-living organisms (the *Carpediemonas*-like Organisms or CLOs) that diverged basally to the parasitic diplomonads (Takashita et. al., 2012). Together the CLOs and diplomonad parasites form the lineage Fornicata, which, in turn, are grouped with other major parasitic groups such as the parabasalids (*e.g.*, *Trichomonas vaginalis*) or anaerobic lineages such as the Preaxostyla in the higher taxonomic ranked Metamonada (Figure 1).

**Figure 1.**
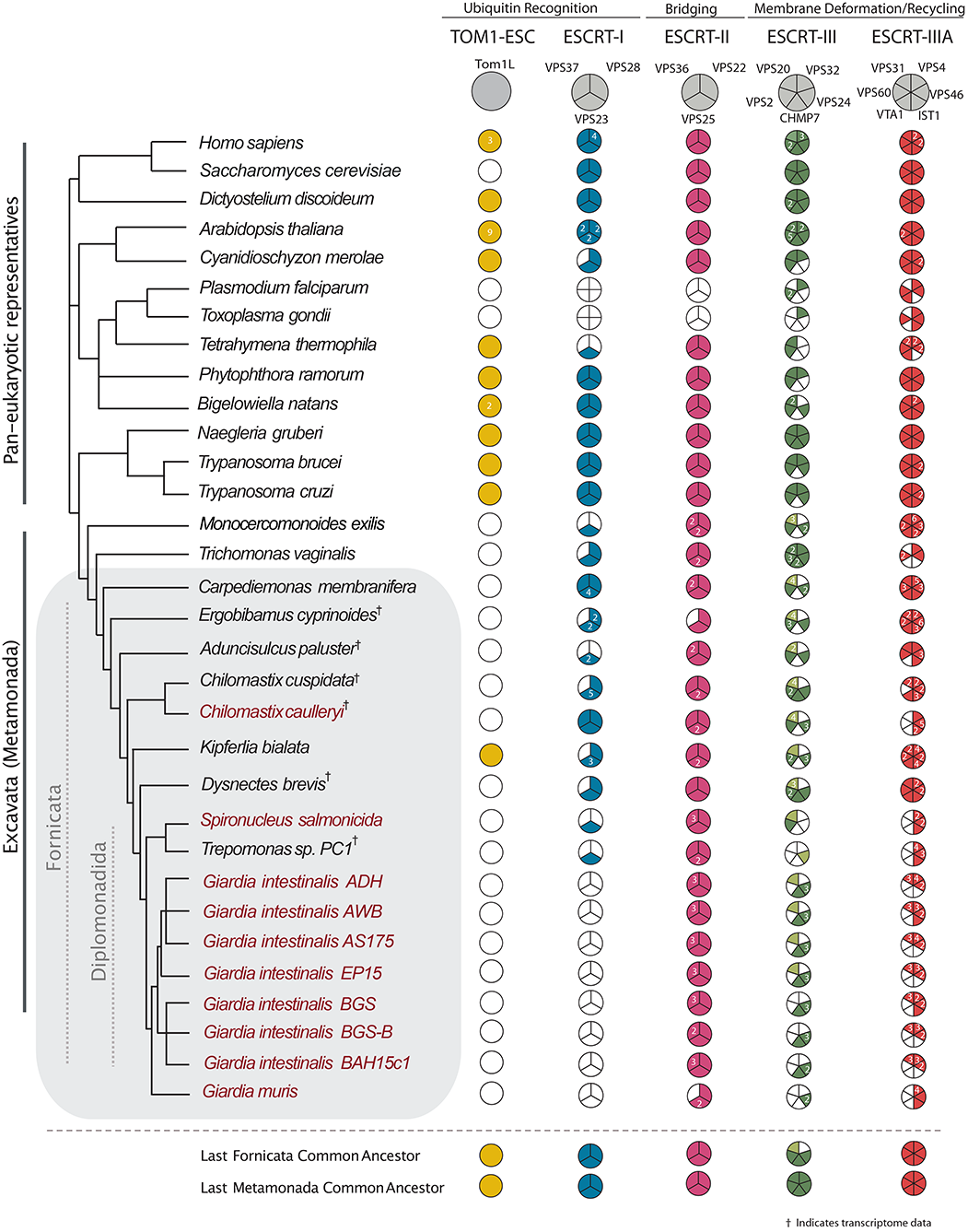
Distribution of ESCRT components within Fornicata. Coulson plot summary depicting ESCRT complement identified in Fornicata genomes and transcriptomes in comparison to pan-eukaryotic representatives. Filled sectors indicate subunits with solidified orthology determined using both comparative genomics and phylogenetics (numbers representing multiple paralogues). Light coloured sectors indicate ambiguous phylogenetic classification but confirmed reciprocal blast orthology. Taxa for which genomes were available and examined are indicated in plain text whereas transcriptomes are indicated with a superscript symbol. Additionally, parasitic lineages are indicated in burgundy. Of important note, only inferences regarding gene presence, not absences, can be made conclusively in the lineages for which only a transcriptome is available.

To date, although ESCRT complement of representative metamonads (*Giardia* included) have been reported, no survey has been done of the entire Fornicata lineage nor other *Giardia intestinalis* assemblages, further raising the important evolutionary question of whether the losses reported in *Giardia* evolved concurrently with parasitism or are a product of gradual evolution that predate its movement into this niche.

Our initial approach using bio-informatics, traces the evolution of the ESCRT system in the Fornicata, finding losses of ESCRT components across the lineage, spanning the transition to parasitism. We also identified several novel components of the ESCRT machinery in *Giardia* and investigated their subcellular localization, revealing ESCRT association at PVs and other locations. Evolutionary modification of the ESCRT complement spans the transition to parasitism in the lineage leading to *Giardia*, and the modified ESCRT machinery acts at more locations than previously understood in this globally important parasite.

## RESULTS

### ESCRT losses in Fornicata are gradual and represent a slow transition leading to parasitism

To understand the extent to which the loss of ESCRT components correlates with parasitism, versus pre-dating it, we investigated the complement encoded in the transcriptomes of free-living *Carpediemonas membranifera* and *Carpediemonas*-like organisms (CLOs) by comparative genomics (Figure 1, Supplementary Table S2). In the case of ESCRT-III and ESCRT-IIIA SNF7 components (VPS20, VPS32, VPS60, VPS2, VPS24, and VPS46) which are themselves homologous, we also used phylogenetic analysis for classification. We took a two-step approach to account for divergent fornicate sequences, first classifying *Carpediemonas membranifera* sequences, and subsequently using these as landmarks to classify the fornicate representatives and then to verify the classification of SNF7 components.

This analysis allowed us to resolve the presence of nearly all SNF7 components with clear clustering with pan-eukaryotic orthologs (Figure 2A, Supplementary Figure S2). The exception was lack of clear VPS32 or VPS20 orthologs in *Carpediemonas.* Instead, multiple VPS20-like proteins were identified. This could imply that one of these protein paralogs may carry out the functions of canonical VPS32 or VPS20 (Figure 2A, Supplementary Figure S2). We do not rule out the possibility that orthologs of VPS32 and VPS20 are present in the *Carpediemonas* gene repertoire which remained unexpressed in standard culturing conditions and, therefore, absent within the assembled transcriptome. Phylogenetic analyses of the identified SNF7 sequences in the remaining CLOs and diplomonads including the five *Giardia intestinalis* isolates further revealed that, similar to *Carpediemonas membranifera*, all VPS20 or VPS32 proteins in all Fornicata lineages have diverged to the extent that no clear clades are resolvable for either VPS20 or VPS32 (Figure 2B, Supplementary Figure S3).

**Figure 2.**
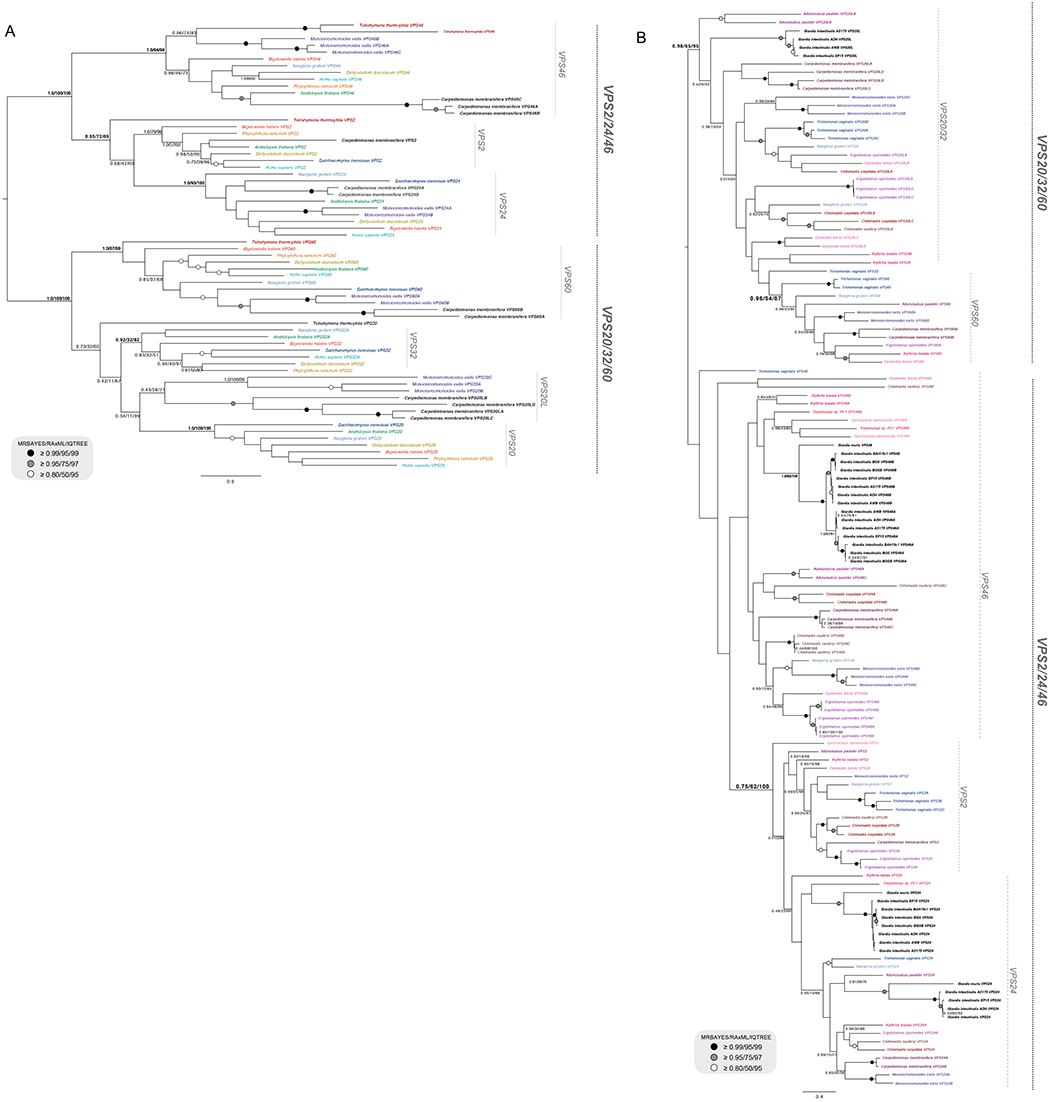
Phylogenetic analysis of ESCRTs in Fornicata. (A) Phylogenetic analyses of the ESCRTIII/IIIA SNF7 families in Fornicata. Identified ESCRTIII/IIIA SNF7 components from the basal *Carpediemonas membranifera* as a landmark representative for Fornicata were subject to phylogenetic classification. Two of the identified SNF7 sequences from *Carpediemonas membranifera* clustered clearly with VPS60 whereas the remainder neither strongly grouped with VPS20 or VPS32 and therefore were determined to be VPS20L proteins in all tree topologies. *Carpediemonas membranifera* was also determined to have VPS2, VPS24 and VP46 with strong backbone clade support for two paralogs of VPS24 (1.0/100/100) and three paralogs of VPS46 (1.0/100/100). (B) A Fornicata-specific tree with well characterized Discoba and metamonad representatives. *Monocercomonoides exilis, Trichomonas vaginalis,* and *Naegleria gruberi* as well as newly characterized sequences from *Carpediemonas membranifera* were used to classify SNF7 components in all CLOs and diplomonads. Similar to *Carpediemonas,* no clear grouping of SNF7 sequences from CLOs within the VPS20 or VPS32 clade was observed and therefore were also classified as VPS20L. Only sequences from *Giardia* AWB, ADH, and EP15 formed a group within this clade and therefore were also determined to be VPS20L. VPS2 family proteins identified in the diplomonads grouped with both VPS24 and VPS46 with duplication event pointing in *Giardia* sp. VPS46 yielding two paralogues, VPS46A and VPS46B. An additional set of SNF7 family proteins from *Giardia* AWB, ADH, and EP15 grouped with excavate and CLO VPS24 proteins therefore were determined to be VPS24-like proteins. However, an additional set of SNF7 proteins from all *Giardia* lineages formed a separate sister clade and therefore also termed to be VPS24. Trees were rooted between the VPS20/32/60 and VPS2/24/46 as previously determined by Leung et al., (2008).

We also notably detected a CHMP7 ortholog in several fornicate representatives, including *Giardia* and the free-living *Chilomastix cuspidata* and *Dysnectes brevis*. We further examined these proteins through domain analysis, which revealed that the characteristic C-terminal SNF7 domain normally required for the recruitment of downstream ESCRT-III VPS20 and VPS32 was absent from all identified CHMP7 orthologs. Following the same pattern as the VPS20L protein, this finding implies partial loss of sequence and divergence of the CHMP7 sequences predates the fornicate common ancestor. Overall, our investigation of the free-living fornicate transcriptomes in direct comparison with the parasitic diplomonads and various isolates of *Giardia* has been useful in retracing the timepoints and instances of ESCRT sequence divergence.

### Losses correlating with parasitism and inter-strain variation

Focusing more specifically on parasitic lineages, including five *Giardia* isolates, the fish parasite *Spironucleus salmonicida,* and the secondarily free-living *Trepomonas* sp. PC1, shows additional losses when compared to their free-living relatives (Figure 1). Within the ESCRT-III machinery, we were unable to classify any of the identified SNF7 proteins as canonical VPS2 proteins in either *Giardia* or *Trepomonas* sp. (Figure 2B, Supplementary Figure S7). Instead, phylogenetic classification pointed towards homology to VPS24 and therefore these proteins were termed VPS24-like (VPS24L) proteins (Figure 2B, Supplementary Figure S7). Additionally, the coincident loss in all diplomonads of VTA1 and VPS60 (Figure 2A) which interact to regulate VPS4 oligomerization hints at the dispensability of the ESCRTIII-A components and that alternative factors, potential paralogs of the unidentified components, may be at play to carry out these functions (Yang et. al., 2012). Other losses common to all diplomonads include ESCRT-I VPS37 and VPS28 that are not only absent in *Giardia* but also *S. salmonicida* and *Trepomonas* sp. PC1. These are indicative of adaptive genome streamlining that likely occurred in the Last Diplomonad Common Ancestor (Figure 1). By contrast, although greater streamlining has occurred in the diplomonads with respect to other fornicates, presence of VPS23 in *Trepomonas* sp. PC1 still hints at the capacity of these lineages to form canonical multivesicular bodies.

We also observed unanticipated differences between the two human infecting assemblages, A and B, at the protein complement level. Assemblage A isolates, AWB and ADH possess two VPS24 paralogues, with one clustering with other canonical VPS24 orthologs from other excavates, the other forming a clearly separate clade, here termed VPS24L (Figure 2B and Supplementary Figure S7). Additionally, we failed to identify any orthologs of VPS20L proteins in assemblage B isolates, BGS and BGS-B. Lastly, we find a similar encoded ESCRT repertoire between the assemblage A, and EP15 strains, as well as phylogenetic clustering of the EP15 sequences with ADH and AWB (Figure 1, Figure 2B), consistent with a proposed closer relationship of these strains to one another than to assemblage B.

Previous work analyzing only the *Giardia intestinalis* AWB ESCRT machinery reported absences in various components such as ESCRT-II VPS36, ESCRT-III CHMP7, and ESCRT-IIIA subunits (Dutta et. al., 2015, Saha et. al., 2018). Here we show these to be present but were not previously detected probably due to high sequence divergence and the lack of the currently available genomes and or transcriptomes from free-living relatives of *Giardia* (Figure 1 and Supplementary Table S2).

### Localization of Giardia ESCRT-II VPS25 and newly identified ESCRT-II VPS36 at peripheral vacuoles

Previous molecular cell biological analyses of ESCRTs in *Giardia* have been limited to highly conserved ESCRT-III and ESCRT-IIIA components (Saha et. al., 2018). The bioinformatic identification of multiple newly described ESCRT components, particularly some with unclear phylogenetic affinity (*e.g.,* VPS20L) make attractive targets for a molecular cell biological approach.

We began by characterizing the ESCRT II component VPS25, which had been consistently identified in previous phylogenetic analyses but never localized. Past work from Giardia on ESCRT III ESCRT components (Saha 2018) and assuming straight forward functional homology from model systems, the simple prediction is for Vps25 to associate with PVs. Immunofluorescence assays of standalone staining in transgenic trophozoites expressing *Gi*VPS25 C-terminally HA-epitope-tagged reporters (*Gi*VPS25-HA), revealed an accumulation in the cell periphery and a punctate cytosolic pattern (Figure 3A and Supplementary Video 1, Figure S4 panel I). Signal overlap analyses on cells (N≥15) labelled for *Gi*VPS25-HA and incubated with the endocytic fluorescent fluid phase marker Dextran coupled to Texas Red (Dextran-TxR) (Figure 3B and Supplementary Video 2, Figure S5 panel I) support partial VPS25 association of VPS25 to PVs (Figure 3B, panels II and III). The signal for Vps25 seemed widespread, but punctate throughout the cell, suggestive of multiple locations. By contrast, the Dextran signal was clearly restricted to the cell periphery, consistent with its denoting PVs. Consistent with these observations the Pearson co-efficient describing overall signal overlap was low, as was the Mander’s co-efficient 1, quantifying the degree of Vps25 overlap with Dextran. However, Mander’s co-efficient 2, describing the degree of Dextran overlap with Vps25 was high as was the Costes value, giving us confidence in our results. Overall, the observations suggest that Vps25 localizes to the PV, consistent with past reports of other ESCRT components functioning at this organellar system. However, it also suggested that Vps25 is found at other organelles within Giardia.

**Figure 3.**
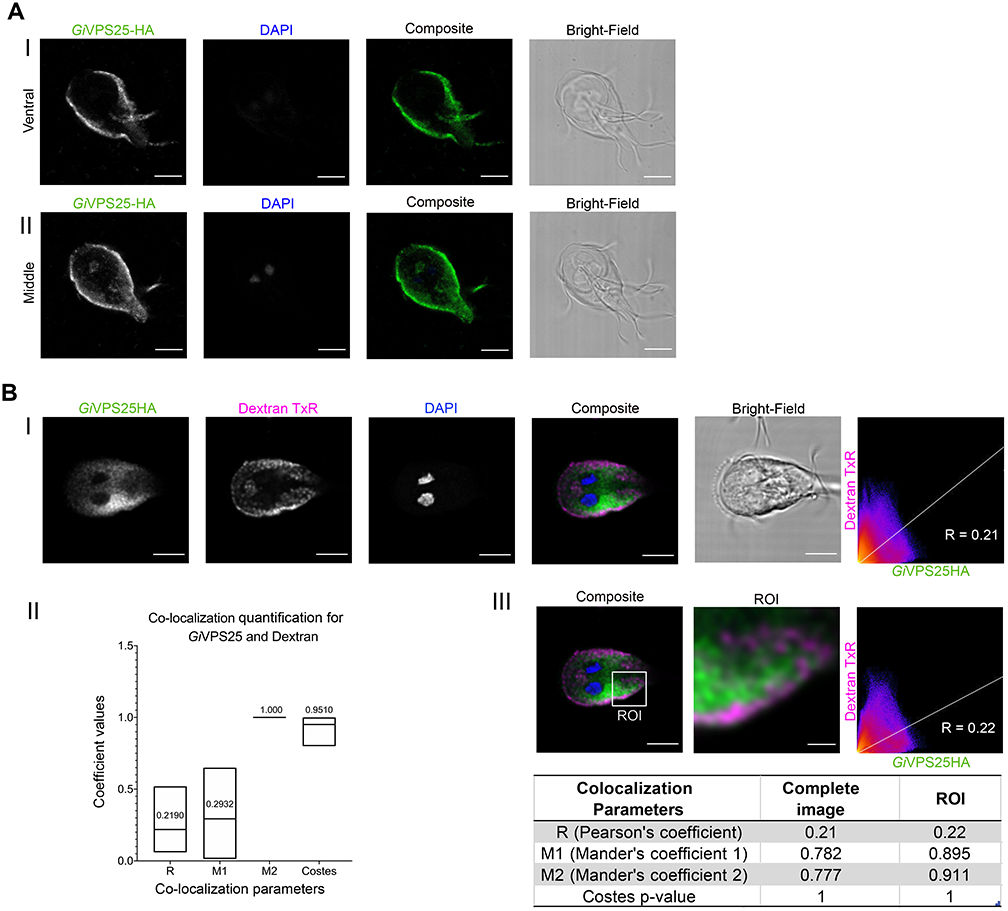
Characterization of *Gi* VPS25-HA subcellular location. (A) Trophozoite cell periphery and cytosol. (I) Ventral and (II) middle optical slice of transgenic *Giardia* trophozoite expressing epitope-tagged *Gi*VPS25-HA (green). All images were obtained using Confocal Laser Scanning Microscopy. All scale bars: 5 μm. (B) PVs. (I) Immunofluorescence-assay of transgenic *Giardia* trophozoite labelled for epitope-tagged *Gi*VPS25-HA (green) and Dextran-TexasRed (magenta). (II) Distribution of co-localization parameters for *Gi*VPS25-HA and Dextran-TexasRed labeling from ≥15 analysed cells. Mean values for each parameter are indicated. (III) Signal overlap analysis and co-localization coefficients calculated for all slices of the sample either for the whole cell or ROI. Scale bars: composite 5 μm and ROI 1 μm. All images were obtained using Confocal Laser Scanning Microscopy.

We proceeded to characterize one of the putative ESCRT components newly identified in our bioinformatic analysis, *Gi*VPS36, hereafter referred to as *Gi*VPS36A (Supplementary Table S2). A molecular cell biological approach here is particularly informative, given that none of the three *Gi*VPS36 paralogues were identified as possessing a *bone fide* GLUE domain. In model systems, this functional module mediates interactions between the ESCRT-I and ESCRT-III sub-complexes (Gill et. al., 2007), which are in turn necessary for ubiquitin-dependent initiation of ILV biogenesis. Instead *Giardia* VPS36 paralogues possess an N-terminal PH domain (Supplementary Table 2), raising questions of functional homology of this component with that of other model organisms. We chose to test localization of *Gi*VPS36A, as this was promptly identified by homology searching, and thus likely to be the least divergent in function. As with VPS25, a localization pattern associated to the cell periphery and punctate cytosolic foci was observed (Figure 4A and Supplementary Video 4, Figure S4 panel II). Signal overlap analyses on cells labelled for *Gi*VPS36A-HA and incubated with Dextran-TxR support partial VPS36A association to PVs (Figure 4B panels II and III and Supplementary Video 5). The Mander’s co-efficients again suggested PV localization, as well as localization to other organelles, an avenue pursued below.

**Figure 4.**
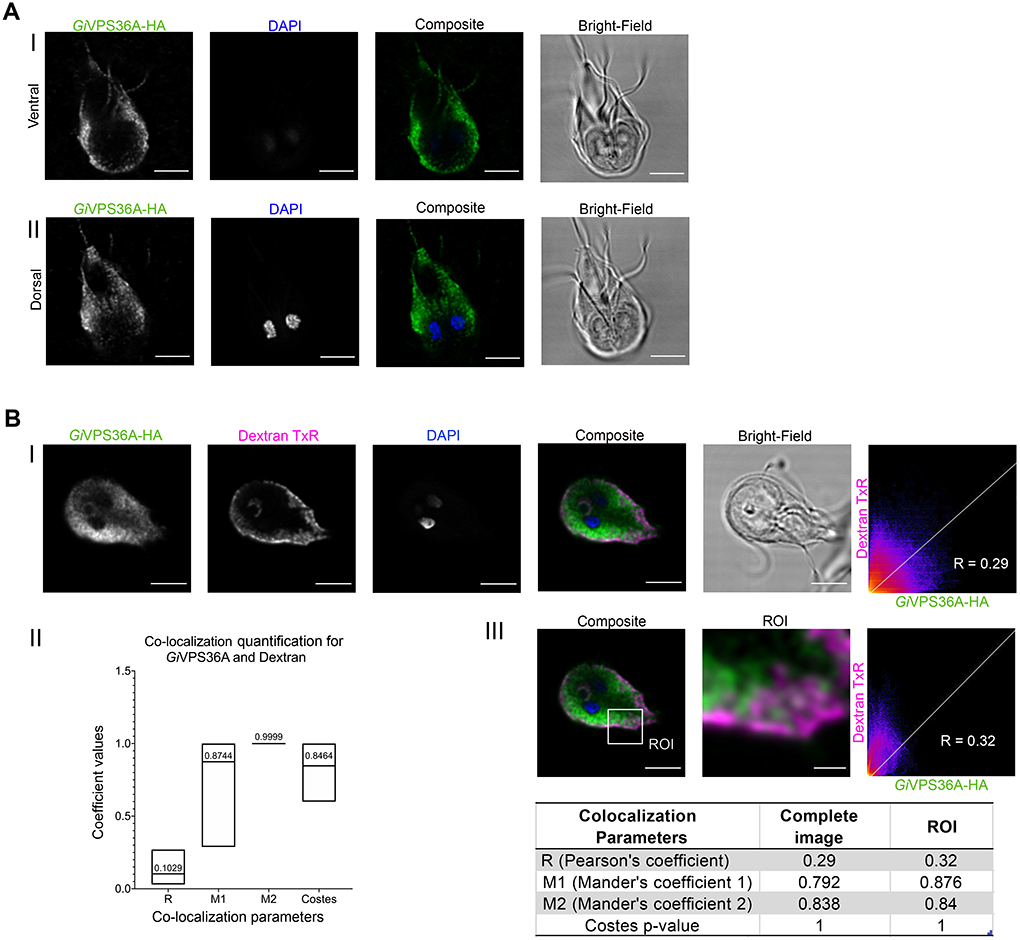
Characterization of *Gi* VPS36A-HA subcellular location. (A) Cell periphery and cytosol. (I) Ventral and (II) dorsal optical slices of transgenic *Giardia* trophozoite expressing *Gi*VPS36A-HA. All images were obtained using Confocal Laser Scanning Microscopy. All scale bars: 5 μm. (B) PVs. (I) Immunofluorescence-assay of transgenic *Giardia* trophozoites labelled for *Gi*VPS36-HA (green) after incubation with Dextran-TxR (magenta). (II) Distribution of co-localization parameters for *Gi*VPS36-HA and Dextran-TexasRed labeling from ≥15 analysed cells. Mean values for each parameter are indicated. (III) Signal overlap analysis and co-localization coefficients calculated for all slices of the sample either for the whole cell or ROI. Scale bars: composite 5 μm and ROI 1 μm. All images were obtained using Confocal Laser Scanning Microscopy.

### Characterization of ESCRT-III VPS20L and ESCRT-II components at the Endoplasmic Reticulum

The newly identified ESCRT-III VPS20L protein family was phylogenetically unresolved in our analyses and the *G. intestinalis* AWB sequence relatively divergent. Both the novelty and divergence of this protein prompted us to investigate this protein further. *Gi*VPS20L was expressed as an N-terminally epitope-tagged reporter (HA-*Gi*VPS20L) and detected by immunofluorescence localization assay (Figure 5A and Supplementary Video 7, Figure S4 panel III) where we observed punctate and dispersed cytosolic localization, previously seen with *Gi*VPS25-HA and *Gi*VPS36A-HA and reminiscent of ER association of cytosolic components (Faso et al., 2013).

**Figure 5.**
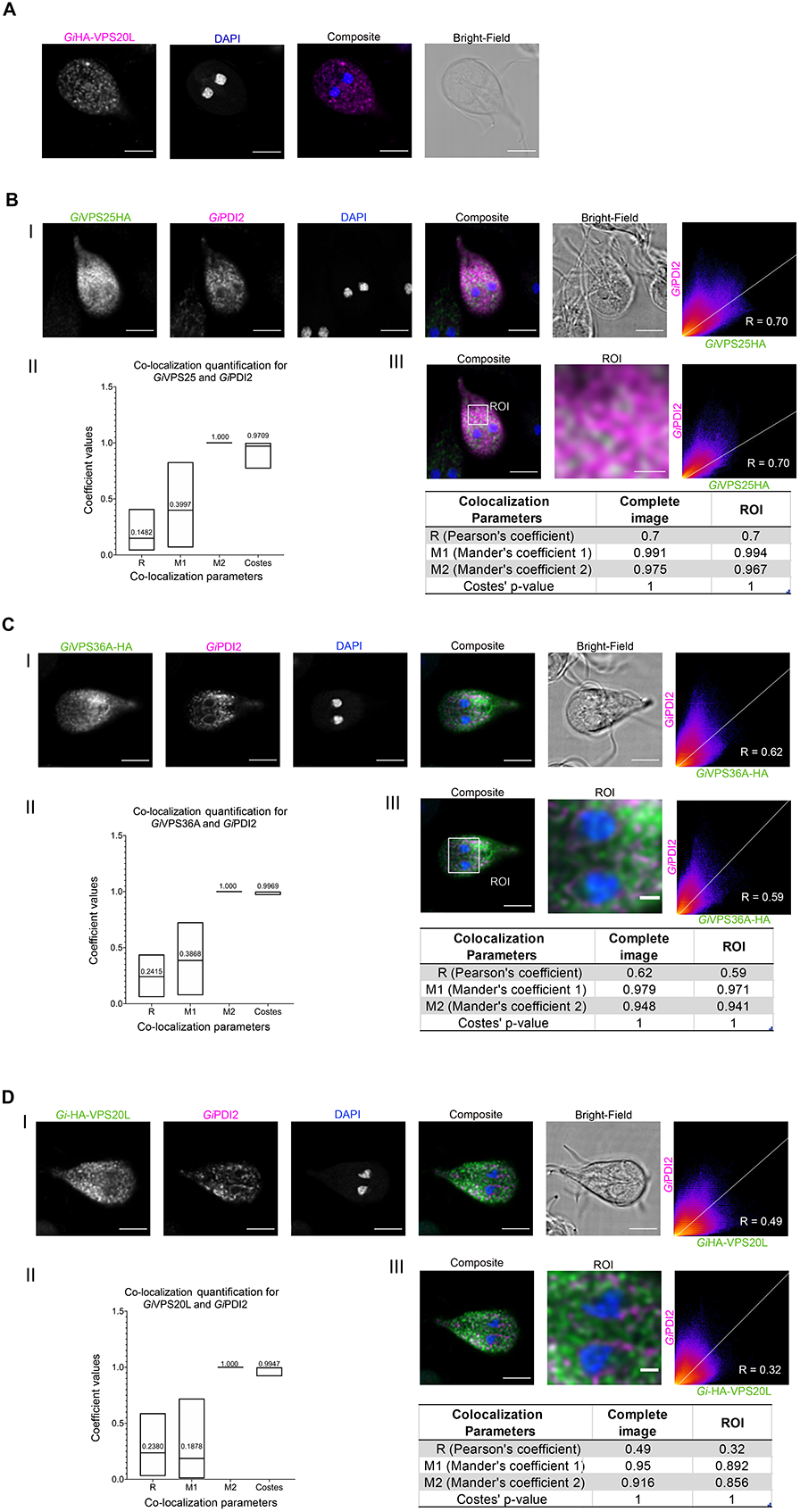
Co-labelling of *Gi*VPS25-HA, *Gi*VPS36A-HA and *Gi*HA-VPS20L with ER membrane marker *Gi*PDI2. (A) HA-*Gi*VPS20L is found in the cytosol and punctate structures. Scale bars: 5 μm. (B-D) Panels I: Co-labelling of PDI2 (magenta) in cells expressing either (B) *Gi*VPS25-HA (green), (C) *Gi*VPS36A-HA (green) or (D) *Gi*HA*-* VPS20L (green). (B-D) Panels II: Mean values from ≥15 analysed cells for each parameter are indicated. (B-D) Panels III: Signal overlap analysis and co-localization coefficients calculated for all slices of the sample either for the whole cell or ROI. Scale bars: composite 5 μm and ROI 1 μm. All images were obtained using Confocal Laser Scanning Microscopy.

This observation, along with the potential for other organellar localization suggested for VPS25 and Vps36, prompted us to investigate whether all three proteins might be ER-associated. To do this, we proceeded with signal overlap analyses of cells (N≥15) co-labelled for each epitope-tagged reporter in combination with the ER membrane marker *Gi*PDI2 (Stefanic et. al., 2006; figures 5B-D). The data shows ESCRT proteins VPS25 (Supplementary Video 3), VPS36A (Supplementary Video 6) and VPS20L (Supplementary Video 8) partially associate to the ER (Figure 5B-D panels II and III). We interpret the low M1 co-efficients (measuring the respective ESCRT components overlap with PDI) but high M2, as most consistent with the ESCRT proteins localized to ER as well as other cellular locations, eg. PVs.

The observation of Vps25 being in ostensibly the same locations at Vps36 and Vps20, at PVs and ER respectively, leads to a prediction that it should show overlap in localization with these two proteins. This was assessed by developing and investigating dually-transgenic *Giardia* lines expressing *Gi*VPS25-HA in combination with either *Gi*VPS36-V5 (Figure 6A panel I) or V5-*Gi*VPS20L (Figure 6B panel I). Based on the signal overlap analysis of co-labelled cells (≥15), there is significant signal overlap in subcellular location for both *Gi*VPS25HA and *Gi*VPS36A-V5 (Figure 6A panels II and III) and *Gi*VPS25-HA and *Gi*V5-VPS20L (Figure 6B panels II and III) in co-expressing cells. Notably, the Pearson co-efficients and both Manders’ coefficients are substantially higher for ESCRT component overlap (Figure 6) than observed for the previous co-localizations against organellar markers (Figures 3,4,5), suggesting we are capturing a consistent picture of a multi-faceted cellular ESCRT localization.

**Figure 6.**
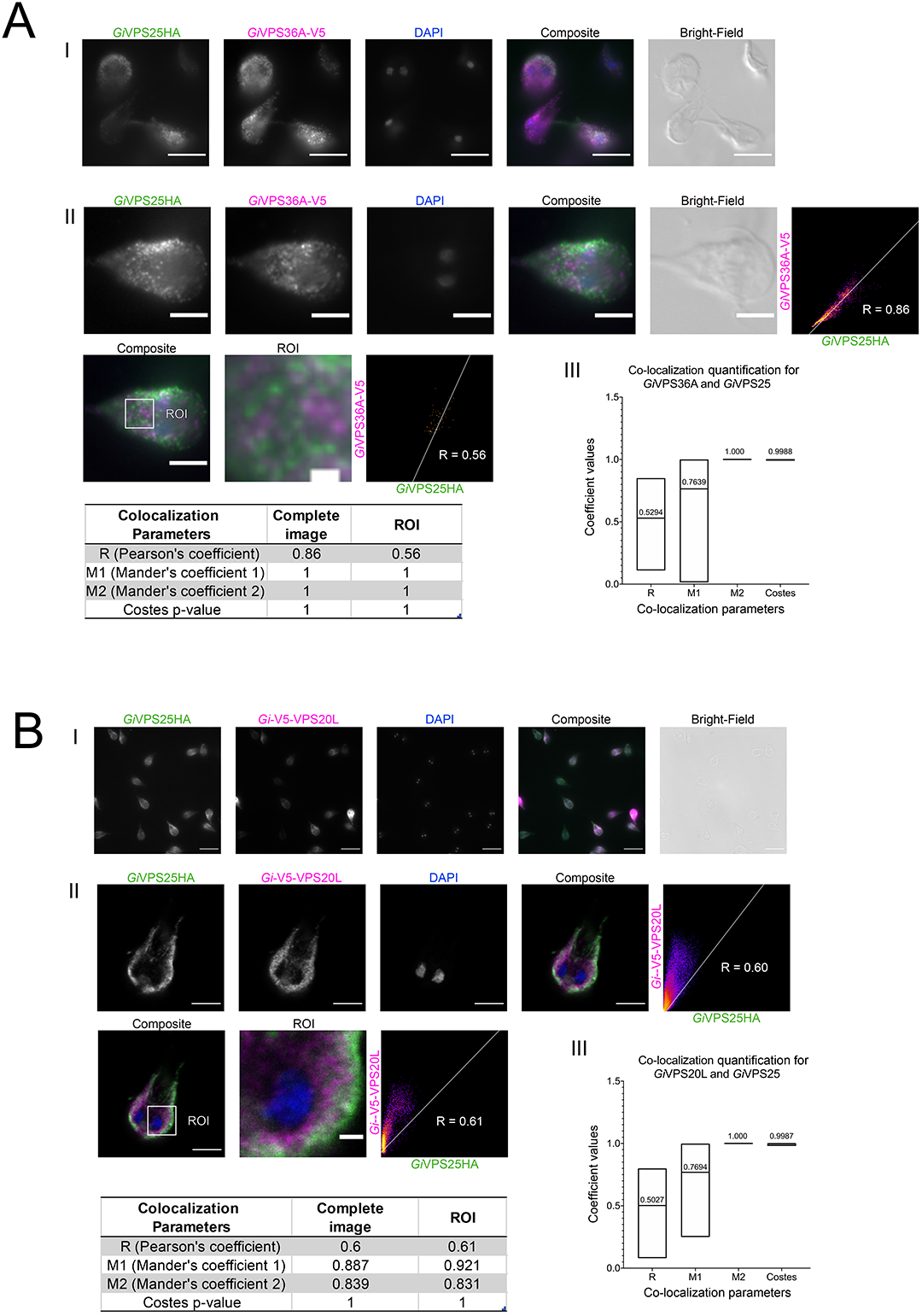
Co-expression of epitope-tagged *Gi*VPS25 with either *Gi*VPS20L or *Gi*VPS36. Microscopy analysis of cells co-expressing *Gi*VPS25HA (green) with either (A) *Gi*VPS36A-V5 (magenta) or (B) *Gi*V5-VPS20L (magenta). Panels I: representative cell images and percentage of co-labeling. Panels II: Signal overlap analysis in both whole cells and regions of interest (ROI). Scale bars: (I) 10 μm, (II whole cell) 5 μm and (II-ROI) 1 μm. Panels III: Mean values from ≥15 analysed cells for each parameter are indicated.

### Evolutionary and protein analyses of the newly identified Giardia ESCRT-III CHMP7 reveal unsuspected ancient origins and a novel ER-Mitosomal interaction

Perhaps the most surprising finding from the comparative genomics analysis was the identification of CHMP7 homologues in multiple Fornicata representatives, despite it being frequently not identified in many genomes across eukaryotes (Figure 1). Fornicate CHMP7 proteins were also highly divergent, missing the C-terminal domain in both *Giardia* and the CLO orthologs.

CHMP7 is currently proposed as being derived from a pre-LECA fusion of two SNF7 domains (Horii et al., 2006). In order to first validate the putative CHMP7 candidates as not being divergent in-paralogs of SNF7, we undertook a combined phylogenetic and structural homology approach. HHPRED and iTASSER analyses of the *Giardia* CHMP7 showed a lack of a predicted C-terminal SNF7 domain. Surprisingly, they also showed sequence and structural homology of the remainder, *i.e.*, the N-terminus of this protein, to the ESCRT-II VPS25 (Figure 7, Supplementary Table 3) rather than to SNF7.

**Figure 7.**
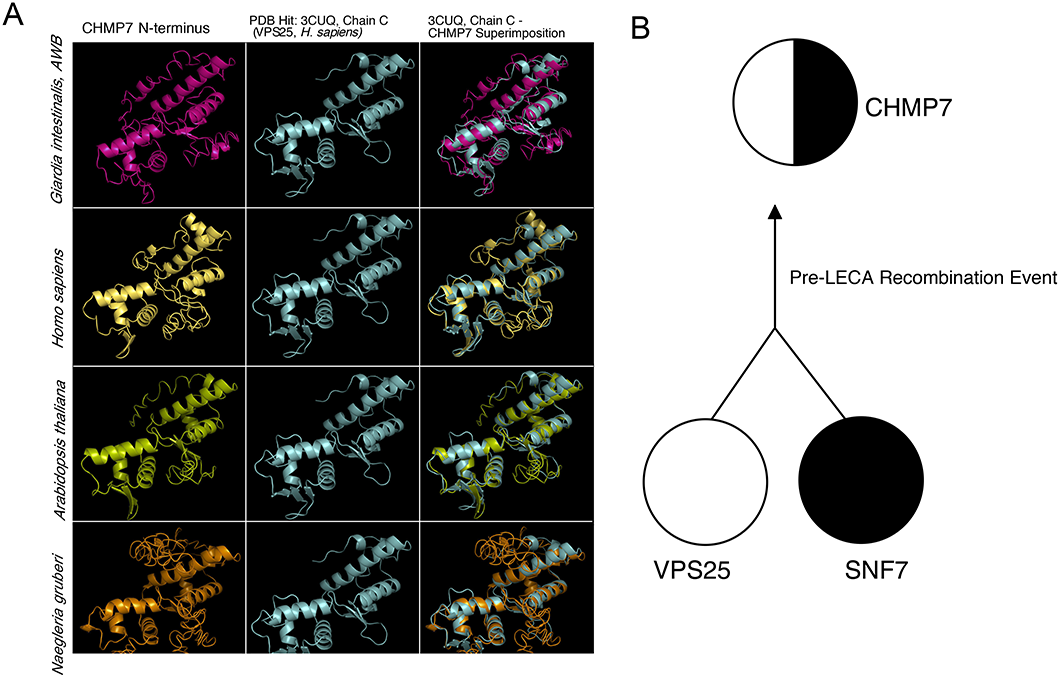
*Ab initio* homology based structural analysis of the CHMP7 N-terminus. Homology-based protein structural analysis of the CHMP7 N-terminus from various pan-eukaryotic representatives carried out using iTASSER *ab initio* structural prediction program where considerable structural similarity between the ESCRTII-VPS25 and CHMP7 N-termini. (B) Proposed evolution as determined by homology searching, structural analyses, and phylogenetic analysis (Supplementary Figures S8-S11) of the pan-eukaryotic CHMP7 protein prior to the last eukaryotic common ancestor which consisted of an evolutionary fusion event between a pre-LECA ESCRTII-VPS25 and ESCRTIII/IIIA-SNF7 progenitor protein.

Analyses with selected CHMP7 N-termini from several representatives of other eukaryotic supergroups confirmed this homology assessment (Figure 7A, Supplementary Table 3). The identity of the fornicate proteins as CHMP7 and not as in-paralogues of VPS25 or SNF7 was also confirmed through our phylogenetic analyses (Figures S9, S11). Our collective structural prediction and phylogenetic findings suggest that a duplication event followed by a fusion event between the VPS20/32 SNF7 and VPS25 had occurred prior to the last eukaryotic common ancestor but subsequent to eukaryogenesis from the presumed Asgard archaeal ancestor.

CHMP7 has been shown to have a variety of functions beyond the endocytic pathway in mammalian or yeast model cell systems (Vietri et. al., 2015, Bauer et. al. 2015, Gu et. al., 2017). Therefore, following the identification of this protein in *Giardia* we aimed to investigate its role in the endomembrane system as well as its relation to the remainder of ESCRTs in this parasite. Based on *Gi*CHMP7’s similarity to VPS25 and lack of a SNF7 domain, we expected similar localization and protein interaction patterns as ESCRT-II components, specifically at the PVs and at the ER. However, our immunofluorescence assay analyses with an N-terminally epitope-tagged *Gi*CHMP7 reporter (HA- *Gi*CHMP7) yielded a distinct localization pattern strongly reminiscent of ER labelling, with no obvious indication of PV association (Figure 8A, Figure S4 panel IV and Supplementary Video 9). As done for *Gi*VPS25-HA, *Gi*VPS36A-HA and *Gi*HA-VPS20L, HA-*Gi*CHMP7 cells were co-labelled for *Gi*PDI2 and a signal overlap analysis was performed (N≥15 cells), showing that HA-*Gi*CHMP7 is partially ER-associated, particularly taking M2 and the Costes values into account (Figure 8B panels II and III and Supplementary Video 10).

**Figure 8.**
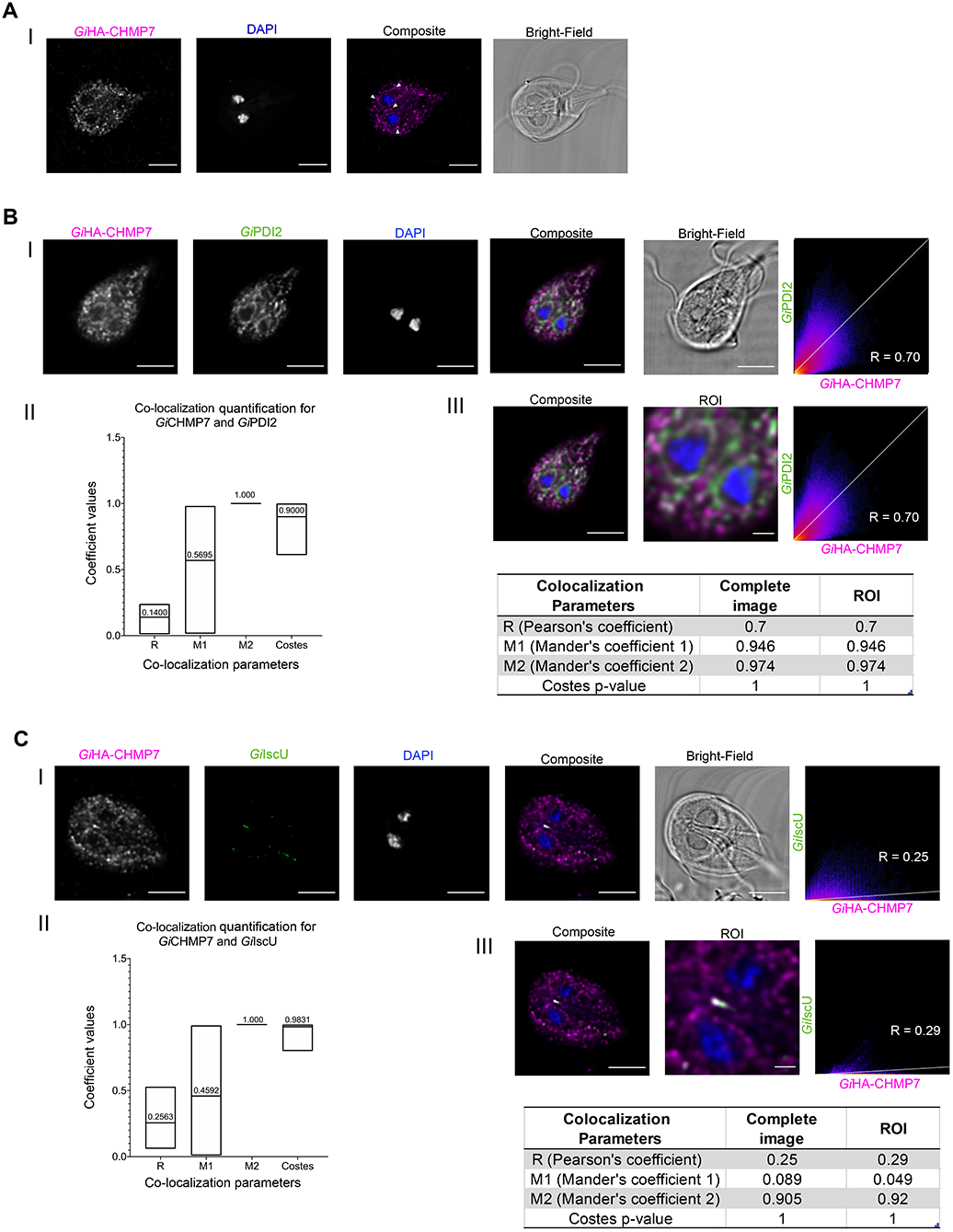
Characterization of *Gi* CHMP7 subcellular location. (A) Immunofluorescence assays of HA-*Gi*CHMP7-expressing cells yield a diffused punctate pattern with elements of perinuclear ER staining (arrowhead). Scale bars: 5 μm. (B) (I) Co-labelling of HA-*Gi*CHMP7 (magenta)-expressing cells with *Gi*PDI2 (green). (II) Distribution of co-localization parameters for *Gi*CHMP7-HA and *Gi*PDI2 labeling from ≥15 analysed cells. Mean values for each parameter are indicated. (III) Signal overlap analysis for all slices of the sample either for the whole cell or ROI. (C) HA-*Gi*CHMP7 is associated to *Giardia* mitosomes. (I) Co-labelling of HA-*Gi*CHMP7 (magenta)-expressing cells with *Gi*IscU (green). (II) Distribution of co-localization parameters for *Gi*CHMP7-HA and *Gi*IscU labeling from ≥15 analysed cells. Mean values for each parameter are indicated. (III) Signal overlap analysis for all slices of the sample either for the whole cell or ROI. Scale bar: composite 5 μm and ROI 1 μm. All images were obtained using Confocal Laser Scanning Microscopy.

Surprisingly, we repeatedly detected CHMP7 signal in compartments consistent with the location of central mitosome complexes (CMC) (Regoes et al., 2005; Figure 8C and Supplementary Video 11). To test this, we co-labelled HA-*Gi*CHMP7-expressing cells with antibodies directed against iron-sulfur cluster assembly component *Gi*IscU to detect mitosomes (Rout et al., 2016; Figure 8C, panel I). We measured significant signal overlap limited to the CMC with *Gi*CHMP7 and *Gi*IscU-derived labels, with the low M2 denoting CHMP7 presence at multiple cellular locales, but very high M2 values indicating strong overlap with the IscU signal (Figure 8C, panels II and III).

## DISCUSSION

*Giardia intestinalis* remains a cause of substantial heath burden world-wide and its divergent cellular and genomic features an enigma from an evolutionary perspective. Our work has specifically addressed the reduced endomembrane system observed in *Giardia intestinalis*, focusing on the ESCRT protein machinery from an evolutionary and molecular cell biological perspective. We show that the reduced ESCRT complement is the product of an evolutionary process that spans the shift from free-living to a parasitic state and includes *Giardia* assemblage-specific losses. We also report on previously unidentified ESCRT machinery and unidentified sites of ESCRT location in *Giardia*, opening novel avenues for investigation.

### Gradual reductive evolution of ESCRTs and MVBs in the Fornicata

Based on the lifestyles of the basally paraphyletic assemblage of CLOs, including *Carpediemonas*, the ancestor of Fornicata was likely a free-living anaerobic flagellate (Leger et. al., 2017). In these conditions, membrane trafficking machinery would be expected to play essential roles in phagotrophy, material exchange, osmoregulation, and intracellular homeostasis. From our analysis, this ancestor appears to have possessed a relatively complete complement of ESCRT machinery as compared with the deduced complement in the LECA. That said, there were likely some component losses that had already taken place (Figure 9), including the CHMP7 SNF7 C-terminus normally required for association with the ESCRT-III VPS32. While it is technically possible that “true” orthologs of these proteins may be encoded in the not-yet sequenced genomes of CLOs, given that the pattern remains consistent across 14 different sampling points, it is much more likely for an ancestral loss to have occurred in the ancestor of fornicates, rather than multiple instances of unexpressed protein or independent losses.

**Figure 9.**
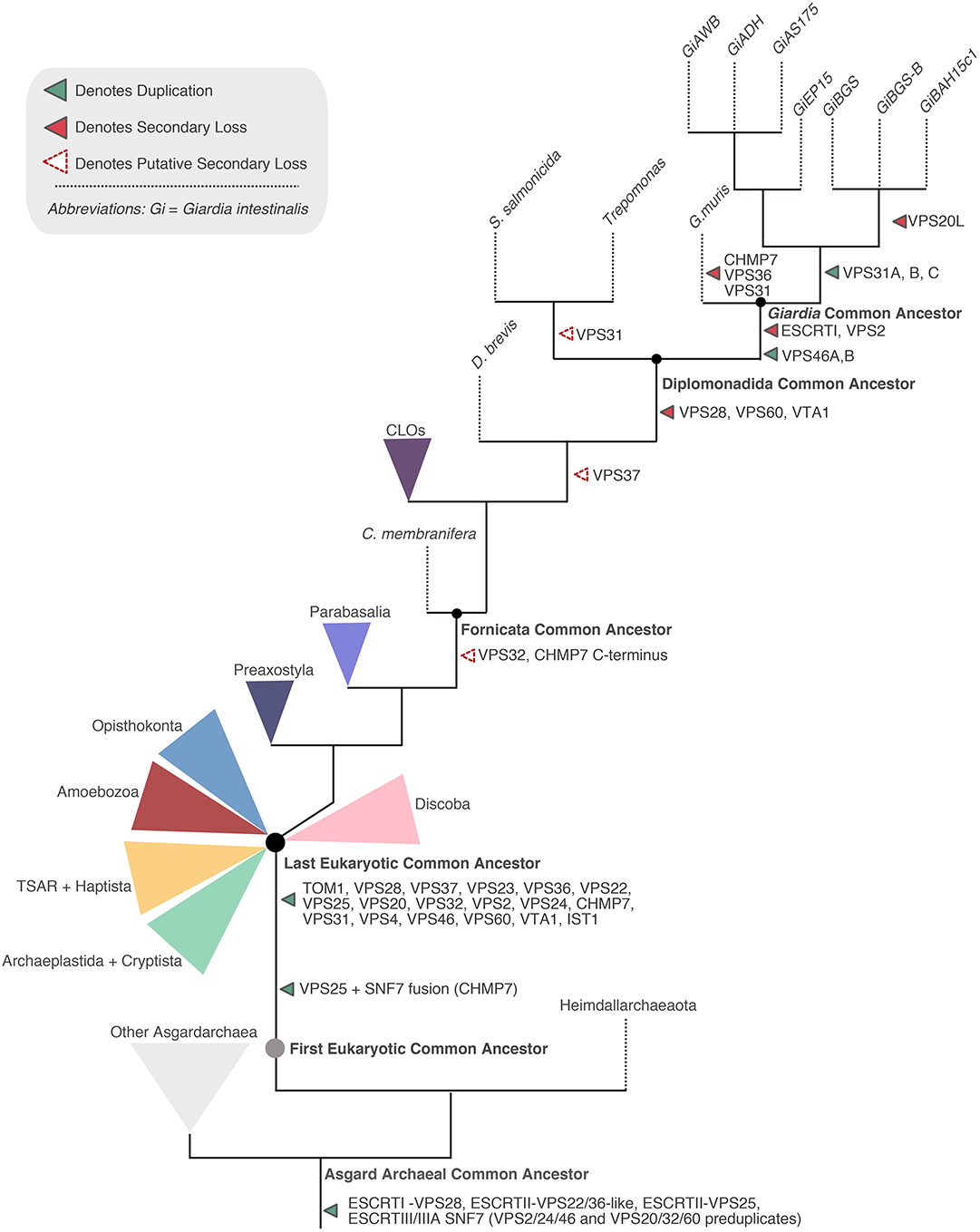
Proposed ESCRT evolution in Fornicata. Progenitor ESCRT complexes are present in Asgard archaea and duplications into the specific subunits is inferred to have occurred between the First Eukaryotic Common Ancestor and the Last Eukaryotic Common Ancestor which possessed a full complement of the ESCRT subunits. Proposed ESCRT losses in Fornicata inferred previously only using *Giardia intestinalis* (Leung et al., 2008; Saha et al. 2018) are transient with some losses potentially predating the Last Fornicata Common Ancestor. The most prominent of this being loss in CHMP7 c-terminus SNF7 domain and a canonical VPS32. Examination of diplomonad lineages, specifically genomic data, increases our confidence in additional losses also occurring with progression into parasitism most notable within the ESCRTI machinery with complete loss occurring in the *Giardia* common ancestor likely associated with a loss in the canonical MVB morphology. Speculative losses indicated as unfilled dotted arrows whereas instances of likely true gene absence depicted as solid filled arrows.

Loss in the SNF7 domain of CHMP7 may functionally relate to the other deduced loss observed in all free-living fornicates, that of a canonical VPS32 protein. By contrast, the transition to parasitism appears to have happened by the time of the diplomonad common ancestor. Concurrent with this are losses of VPS28, VPS60, VTA1 and possibly VPS37 (Figures 1 and 9). These are correlated, though not necessarily causally associated, with this transition. Notably, however, VPS23 is retained in some diplomonads and is characterized by the presence of a UEV domain which is required for interaction with cargo tagged with Ubiquitin for targeted lysosomal degradation. Lineages such as *Tetrahymena*, *Entamoeba*, and *Monocercomonoides* conserving only the VPS23 from ESCRT-I appear to be capable of forming functional (or at least morphologically identifiable) multivesicular bodies (Cole et. al., 2015, Okada et. al., 2006, Karnkowska et. al., 2019). In turn, this allows us to predict that all fornicate lineages possessing ESCRT-I VPS23, including the diplomonads *S. salmonicida* and *Trepomonas* sp. PC1, may also possess bona fide MVBs.

In the common ancestor of *Giardia* itself, we observed loss in all ubiquitin binding components and domains. Collectively, these include TOM1-esc, ESCRT-I, and VPS36 GRAM and NZF domains. We speculate that the observed lack of canonical MVB morphology in *Giardia intestinalis* specifically corresponds to losses within these components. Instead, we propose that the existing repertoire of *Giardia* ESCRT machinery has an altered role at the *Giardia* specific late endolysosomal organelle, the PVs. Notably, we also observed variability between the different *Giardia* genomes in some ESCRT-III and −IIIA components indicating that there is inter-strain variability in the membrane-trafficking complement worthy of further investigation.

PV vesicle-like contents have been recently observed in *Giardia* (Midlej et. al., 2019: Moyano et al., 2019). However, although *Giardia* may secrete non-exosomal and non-MVB-derived extracellular vesicles, the absence of key ESCRT machinery (e.g. TOM1-esc, ESCRT-I, and VPS36 GRAM domain), along with a lack of specific exosomal markers and limited proteomics data (Ma’ayeh et al., 2017; Coelho et al., 2018) keeps the status of PV-associated vesicles in some doubt. An alternate interpretation is the PVs as a form of reduced and functionally limited MVB-like compartments in a similarly reduced *Giardia* endomembrane system which evolved via a process of merging organelle identity and distribution of endocytic function. Although PVs may not have a direct organellar homologue, it is still meaningful to understand which processes have been distributed to which organelles in this re-organization.

### ESCRT promiscuity at Giardia PVs, ER, and mitosomes

Previous investigations of the *Giardia* ESCRT-IIIA components determined a possible role for this complex at the endolysosomal peripheral vacuoles (Dutta et. al., 2015, Saha et. al., 2018). While ESCRT-IIIA components VPS4 and VPS46 are universally conserved in all eukaryotes, ESCRT-II is not (Leung et al., 2008). Therefore, we aimed to investigate the role of this protein complex that is usually required for bridging an existing ESCRT-I and ESCRT-III in the multivesicular body pathway and how *Giardia* may be utilizing it in the absence of ESCRT-I.

The imaging data and signal overlap analyses performed with tagged reporters for both *Gi*VPS25 and *Gi*VPS36 and fluorescent dextran as a soluble PV lumen marker support a PV association for both ESCRT components. The link between ESCRTs and the endocytic pathway and PVs is further corroborated by cross-referencing previously published coIP datasets derived from PV-associated endocytic components. This highlights the presence of ESCRT proteins in these PV-centric interactomes (Cernikova et. al., 2020, Zumthor et. al., 2016). Tagged reporters for α and β subunits of AP2 collectively immunoprecipitated ESCRT components *Gi*VPS36B, *Gi*VPS36A, *Gi*VPS4A, *Gi*VPS4B, *Gi*IST1, *Gi*VPS24A and the three *Gi*VPS31 paralogs (Zumthor et. al., 2016). *Giardia*’s first characterized dynamin-related protein pulled down ESCRT-IIIA VPS46B, VPS31A and VPS31C. *Giardia* Clathrin heavy and putative light chains’ interactomes (Zumthor et al., 2016), similar to interactomes for the predicted PH-domain carrying PV-associated *Gi*NECAP1 protein (Cernikova et. al., 2020), include ESCRT-IIIA subfamily components *Gi*VPS4A, *Gi*VPS4B and the three paralogs of *Gi*VPS31. This wealth of previously-reported targeted proteomics data points to a clear association of *Giardia* ESCRT components to PVs, further strengthening these organelles’ status as functionally reduced and non-motile endo-lysosomal compartments. A clear association between ESCRT components and the ER also emerged from our investigations and is in line with reports for ESCRT-III participation in budding vesicles from the ER (Mast et al., 2018) and for CHMP7 deposition at the perinuclear envelope (Olmos et al., 2016).

The most surprising association reported here is between CHMP7 and the mitosomal marker IscU, from which we infer a role for this ESCRT component at mitosomes. Notably this inference is corroborated by the presence of *Gi*CHMP7 and ESCRT-IIIA components *Gi*VPS4B, *Gi*VPS46B and *Gi*VPS31in the interactome of mitosome-localized *Gi*MOMTiP1 protein, a main interacting partner of *Gi*Tom40 (Rout et al., 2016). Recent reports point to novel links between ESCRTs, mitochondrial membranes (Richardson et. al., 2014) and mitophagy (Hammerling et. al., 2017; Zhen et. al., 2017, Anding et. al., 2018). Therefore, although ESCRTs have been associated to mitochondria, to our knowledge this is the first report to show an association to mitochondria-related organelles, representing a novel facet of MRO biology that should be explored in *Giardia* and in other MRO-possessing organisms.

Notably, the co-localization co-efficients that we observed for the various ESCRT components told a consistent, if not entirely straight-forward story. In all cases, we observed low co-efficients for overall signal overlap and degree of overlap between the ESCRT component and with discrete organellar markers, but high overlap between the organellar markers and the component. The overall overlap quantification between ESCRT components, especially Vps25 and Vps20L or Vps36 however, were higher indicating that their signals were consistent. Together this tells a story of ESCRT localization at multiple locations, beyond the PV to the ER and even the mitosome in the case of CHMP7.

### A comprehensive appreciation of ESCRT evolution and distribution in Giardia intestinalis

Definition of *Giardia* ESCRTs subcellular localizations combined with rigorous phylogenetic analyses point to streamlining and loss of canonical MVB morphology within Fornicata, mirrored by the selective loss of ESCRT-I, of which *Giardia* is a notable example. In the *Giardia* lineage, we observe duplications in the ESCRT-IIIA machinery with paralogs (Figure 9) which may compensate for ESCRT-I and −III losses while, in combination with remaining ESCRT components, still functioning at PVs. We further observe deep adaptation in *Giardia*’s ESCRT pathway by ESCRT-III components such as the CHMP7 apparently not associating within the endocytic pathway as first proposed (Horii et al., 2006).

In comparison to ESCRT machinery in characterized model organisms (Figure S1), we observe localization of ESCRT-II together with previously analyzed ESCRT-IIIA VPS46 and VPS4 components in close proximity to PVs and ER, while ESCRT-III CHMP7 and VPS20L seem to localize almost exclusively in regions overlapping with the ER, with additional unknown roles for ESCRT-III CHMP7 at mitosomes.

*Giardia* ESCRT-III’s association to the ER and to mitosomes presents a complex landscape of novel membrane remodelling sites while maintaining PVs as reduced and simplified MVB-like compartments mostly by the action of ESCRT-II and ESCRT-IIIA subunits. Our collective data sheds light on a potential mode of action for ESCRT-II and ESCRT-IIIA at the PV membranes. We speculate that these subunits likely associate to the PV outer membrane from a cytosolic pool and perform membrane deformation, as characteristic of other eukaryotic ESCRT subunits. Contacts sites between ER and PVs have been previously documented (Zumthor et. al. 2016) and could additionally be mediated by ESCRTs, allowing protein recycling down the endocytic and secretory pathway.

## CONCLUSIONS

We have traced the evolutionary trajectories of ESCRTs within the Fornicata, observing a slow streamlining of the ESCRT machinery across the transition to parasitism, with losses predating, concurrent with and post-dating. Several groups have recently reported on a broader set of ESCRT functions in the eukaryotic cell than previously understood. In Giardia, ESCRTs have been primarily previously reported at the PV. Additionally, we have shown ESCRT association to other membrane locations such as the ER and mitosome surface, suggesting this machinery may act more extensively at multiple organelles in *Giardia* than expected. Future functional studies should build on this comprehensive report to determine if *Giardia’s* status as a highly-diverged non-canonical eukaryote may be a notable exception when it comes to ESCRT function and complement.

## MATERIALS AND METHODS

### Taxa Studied

The previously-published draft genomes of *Kipferlia bialata* (Tanifuji et al., 2018), genome of *Spironucleus salmonicida* (Xu et al., 2014), transcriptome of *Trepomonas* sp. PC1 (Xu et al., 2016), genome of *Giardia intestinalis* Assemblage AI, isolate AWB (Morrison et al., 2007), genome of *Giardia intestinalis* Assemblage AII, isolate DH (Adam et al., 2013), draft genome of *Giardia intestinalis* Assemblage B, isolate GS (Franzen et al., 2009), genome of *Giardia intestinalis* Assemblage B, isolate GS-B (Adam et al., 2013), and genome of *Giardia intestinalis* Assemblage E, isolate P15 (Jerlström-Hultqvist et al., 2010) were obtained from GiardiaDB and National Centre for Biotechnology Information (NCBI). Latest assemblies were used in each case.

### Translation of Carpediemonas-like organism nucleotide transcriptomes

Nucleotide transcriptomes of *Carpediemonas membranifera* and five *Carpediemonas*-like organisms (CLOs), *Aduncisulcus paluster, Ergobibamus cyprinoides, Dysnectes brevis, Chilomastix cuspidata,* and *Chilomastix caulleryi* were obtained from Dryad Repository (doi 10.5061/dryad.34qd7) (Leger et. al, 2017) and translated using ab initio gene prediction program, GeneMarkS-T under the default parameters (Tang, Lomsadze, Borodovsky, 2015).

### Comparative genomics and homology searching

Query protein sequences for individual subunits from each ESCRT sub-complex from various pan-eukaryotic representatives were obtained and aligned using MUSCLE v3. 8.31 (Edgar, 2004) (Supplementary Table 1). Resulting alignments were used to generate Hidden Markov Models using the hmmbuild option available through the HMMER 3.1.b1 package and HMMer searches into all Fornicata genomes and transcriptomes using the hmmsearch tool with an e-value cutoff set to 0.01 (Eddy, 1998). Non-redundant forward hits were deemed positive if BLASTp reciprocally retrieved the correct ortholog in *Homo sapiens* protein database with an e-value > 0.05 and were two-fold higher in e-value than the next best hit. Reciprocal hits were extracted and sorted using an in-house Perl script.

Additional analyses of hits that failed to retrieve any reciprocal hits were analysed by BLASTp in the NCBI non-redundant database. Additional orthology assessment was carried out using the HHPRED suite for an HMM-HMM profile comparison and predicted secondary structure homology comparison with proteins deposited in the Protein Data Bank (Berman et al., 2008). In order to rule out any false negatives, additional translated nucleotide (tBLASTn) searches were carried out in the Fornicata nuclear scaffolds for components that remained unidentified in HMMER searches. In cases where diplomonad sequences were unidentified due to extreme sequence divergence, identified *Carpediemonas membranifera* and CLO ESCRT orthologs were used to search the diplomonad predicted protein databases by subsequently adding these sequences into the previously generated HMM profile to build a new HMM matrix. Exhaustive BLASTp and tBLASTn analyses were also performed using *Carpediemonas membranifera* and CLO sequences in the nuclear scaffolds of all diplomonads. All Fornicata ESCRT orthologs identified by this method were subject to domain analyses using Conserved Domain Database (CDD) with an e-value cut-off first set at 0.01 and then at 1.0 to detect for any highly diverged domains (https://www.ncbi.nlm.nih.gov). All confirmed hits are listed in Supplementary Table 2.

CHMP7 structural analyses was carried out using HHPRED as described above for select pan-eukaryotic orthologs as well as the *ab initio* structural prediction tool iTASSER for protein threading and secondary structure prediction (Roy et al., 2010). HHPRED results are summarized in Supplementary Table 3.

### Phylogenetic analyses of the ESCRT-III and ESCRT-IIIA SNF7 family proteins

Phylogenetic analyses of the evolutionarily paralogous SNF7 family proteins belonging to ESCRT-III and ESCRT-IIIA sub-complexes was carried out using Bayesian and maximum likelihood approaches (Leung et al., 2008). Identified *Carpediemonas membranifera* ESCRT genes belonging to the SNF7 family were used as a landmark representative (VPS2, VPS24, VPS20, VPS32, VPS46, and VPS60) and were aligned to a pan-eukaryotic backbone containing characterized SNF7 proteins for classification into specific protein families backbone alignment containing pan-eukaryotic sequences as resolved and published by Leung et al using the profile option in MUSCLE v3.8.31 (Leung et al., 2008, Edgar, 2004). Alignments were visualized in Mesquite v3.5 (Maddison and Maddison, 2018) and manually adjusted to remove gaps and regions lacking homology. Upon classification of the *Carpediemonas* sequences, a metamonad-specific phylogenetic analysis was carried out for the classification of identified *Giardia* and diplomonad SNF7 sequences using the same process as described above. An additional set of phylogenetic analysis was repeated using only ESCRT-III and −IIIA components, VPS2, VPS24, and VPS46 specific tree and VPS20, VPS32, and VPS60 specific tree.

Maximum likelihood approaches using non-parametric and ultrafast bootstrapping was performed using RAxML-HPC2 on XSEDE v8.2.10 and IQTREE, respectively (Stamatakis, 2014, Nguyen et al., 2015). For RAxML analyses, protein model testing was performed using ProtTest v3.4.2 (Darriba et al., 2011). In all cases, LG + F + Γ model was used. 100 non-parametric bootstraps with the default tree faster hill climbing method (−f b, −b, −N 100) was carried out. A consensus tree was obtained using Consense program available through the Phylip v3.66 package (http://evolution.genetics.washington.edu/phylip.html) (Felsenstein, 1989). IQTREE best protein model selections were determined using the in-built ModelFinder package (Kalyaanamoorthy et al., 2017). In all cases, LG+F+G4 was determined to be the best-fit model according to the Bayesian Information Criterion. Ultrafast bootstrapping with IQ-TREE v. 1.6.5 were performed using 1000 pseudoreplicates (Nguyen et al., 2015). Bayesian inference was carried out using MRBAYES on XSEDE v3.2.6 with 10 million Markov Chain Monte Carlo generations under a mixed amino acid model with number of gamma rate categories set to 4 (Huelsenbeck and Ronquist, 2001). Sampling frequency was set to occur every 1000 generations and burnin of 0.25 to discard the first 25% of samples from the cold chain. Tree convergence was ensured when average standard deviation of split frequency values fell below 0.01. Random seed value of 12345 was chosen for all phylogenetic analyses. Non-parametric and ultrafast bootstraps obtained from RAxML and IQTREE analyses were overlaid onto the MRBAYES tree topologies with posterior probabilities. RAxML and MrBAYES analyses were performed on CIPRES portal (http://www.phylo.org/index.php) while the IQTREE package v1.6.5 was installed and run locally (Miller et al., 2015). All trees were visualized and rooted in FigTree and annotations were carried out in Adobe Illustrator CS4. All masked and trimmed alignments available upon request.

### Giardia cell culture and transfection

*Giardia intestinalis* strain AWB (clone C6; ATCC catalog number 50803) trophozoites were grown using standard methods as described in Morf et. al. (Morf et al., 2010). Episomally-transfected parasites were obtained via electroporation of the circular pPacV-Integ-based plasmid prepared in *E. coli* as previously (Zumthor et. al., 2016). Transfectants were selected using Puromycin (final conc. 50μgml^−1^; InvivoGen). Transgenic lines were generated and analyzed at least thrice as soon as at least 20 million transgenic cells could be harvested *i.e.* ca. 1.5 weeks post-transfection. Based on microscopy analyses of immunofluorescence assays to detect reporter levels, 85-92% of cells expressed their respective transgene(s) (Supplementary Figure 4).

### Construction of expression vectors

Oligonucleotide sequences for construct generation are listed in Supplementary Table 4. Open reading frames of interest were cloned in the pPacV-Integ vector under control of their putative endogenous promoters. Putative endogenous promoters were derived 150bps upstream of the predicted translation start codon. ORFs were cloned in a modified PAC vector (Wampfler et al., 2014, Zumthor et. al., 2016, and Cernikova et. al., 2020).

### Immunofluorescence Assays

Chemically fixed cells for subcellular recombinant protein localization were prepared as previously described (Konrad, Spycher, and Hehl, 2010). HA-epitope tagged recombinant proteins were detected using a rat-derived monoclonal anti-HA antibody (dilution 1:200, Roche) followed by a secondary anti-rat antibody coupled to AlexaFluor 594 fluorophores (dilution 1:200, Invitrogen). For co-localization experiments with ER or mitosomal markers, samples were incubated with either a mouse-derived anti-*Gi*PDI2 (Stefanic et. al. 2006) or a mouse-derived anti-*Gi*IscU (Rout et. al. 2016) primary antibodies both at a dilution of 1:1000, followed by incubation with anti-mouse antibodies coupled to AlexaFluor 488 fluorophores (dilution 1:200, Invitrogen). For the labelling of the V5 epitope, we used anti-V5 primary antibody (1:1000; Thermofisher) followed by anti-mouse antibodies coupled to AlexaFluor 594 fluorophores (dilution 1:200, Invitrogen). Samples were embedded in Vectashield (VectorLabs) or Prolong Diamond Mounting medium (Invitrogen) containing 4’,6-diamidino-2-phenylindole (DAPI) for nuclear staining.

### Fluid phase marker uptake

Dextran uptake assays were performed as described in (Gaechter et al. 2008; Zumthor et al. 2016) using Dextran 10kDa TexasRed at 2mg/mL (Invitrogen). Immunostaining was performed as described above with the exception of using only 0.05% Triton-X100 (Sigma) in 2% BSA (Sigma) for permeabilization, to prevent leakage and loss of Dextran signal.

### Microscopy and Image Analysis

Imaging was performed in an inverted Confocal Laser Scanning Microscope Leica SP8 using appropriate parameters. Confocal images were subsequently deconvolved using Huygens Professional (https://svi.nl/Huygens-Professional) and analysed using Fiji/ImageJ (Schindelin et al. 2012). For co-localization analysis the coloc2 Fiji/imageJ plugin was used. For this, automatic background subtraction was performed in Fiji/imageJ and 100 Costes iterations were performed (Costes et. al.,2004). Three-dimensional analysis and videos were performed in Imaris version 9.5.0 (Bitplane, AG) (Supplementary Videos 1-11). For statistical analysis of labelling signal overlap between ESCRT subunits and specified markers, a macro was developed in Fiji/ImageJ (Schindelin et al.,2012) (version 1.53d). This script has been made available through supplementary materials (Supplementary File 1). Briefly, each channel was thresholded via WEKA segmentation – a machine learning pipeline (Arganda-Carreras et al., 2017). The derived binary image is used as a mask for signal overlap on ≥15 cells per sample/line using the Fiji plugin coloc2 (Cosets et al., 2004).

## Data Availability

Masked and unmasked protein sequence alignments used for all phylogenetic analyses available upon request.

## ACKNOWLEDGMENTS

All microscopy imaging and image analysis was performed with equipment provided by the Center of Microscopy and Image Analysis (ZMB) at the University of Zurich and the Microscopy Center (MIC) at the University of Bern. We thank members of the Dacks lab, Faso and Hehl groups for insightful discussions.

No conflicts of interest present.

SVP, RS, CF, DSL, AJR, ABH and JBD designed the studies. SVP, RS, EAB, and CDW performed the experiments. SVP, RS, DSL, CF, and JBD carried out data analyses. SVP, RS, CF, and JBD wrote the manuscript. All authors read and approved the final manuscript.

## SUPPLEMENTARY MATERIAL

### Figures

**Figure S1.**
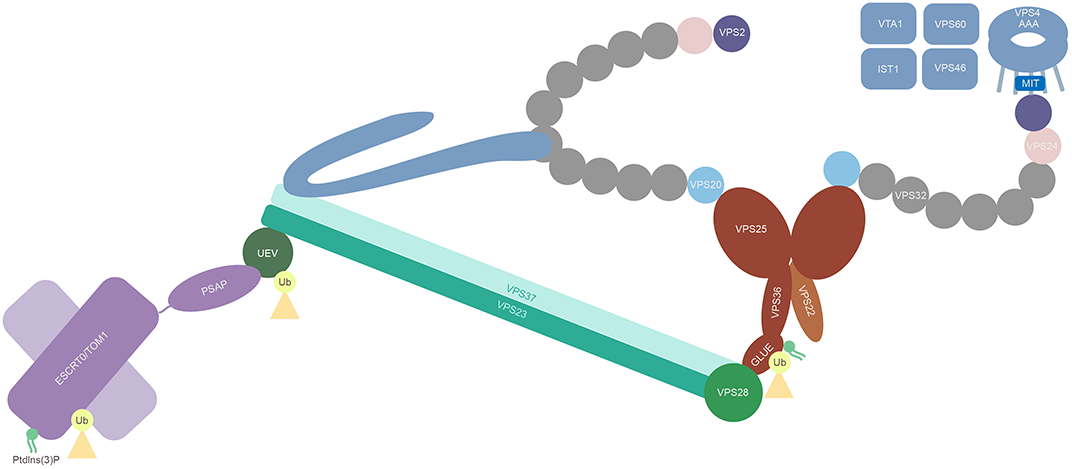
The ESCRT machinery is composed of five sub-complexes each functioning consecutively for recruitment of the downstream subcomplex. The process begins with the recruitment of ESCRT-0 or its analogue TOM1-esc for recognition of tagged Ubiquitin on cargo, and endosomal membrane phospholipids such as phosphatidylinositol 3-phosphate (PtdIns(3)P) upon which the ESCRTI, composed of VPS23, VPS28, and VPS37, is recruited, with its only known role being ubiquitin recognition via its UIM domain (Raiborg and Stenmark, 2009). The assembly of ESCRTI then leads to assembly of the heterotetrameric ESCRTII consisting of VPS36, VPS22, and two copies of VPS25 which also bind to PtdIns(3)P via the FYVE domains (Raiborg and Stenmark, 2009). Finally, this leads to the recruitment of the ESCRTIII machinery, a heteropentameric complex consisting of SNF7 family proteins, VPS20, VPS32, VPS2, VPS24, and CHMP7 (Raiborg and Stenmark, 2009). A filamentous VPS32 polypeptide capped by VPS2 and VPS24 induces ILV formation by constricting the neck of the budding vesicle, a process which is catalysed by the ESCRT-IIIA VPS4, an AAA+ ATPase (Raiborg and Stenmark, 2009). It is also hypothesized that ESCRT-IIIA components such as VPS31 and VPS46 are required for stabilizing the sub-complexes during the budding processes while others are needed for recycling of the complexes back into the cytosol once the process is complete (Odorizzi, 2003). Figure adapted from Stenmark and Raiborg, 2009.

**Figure S2.**
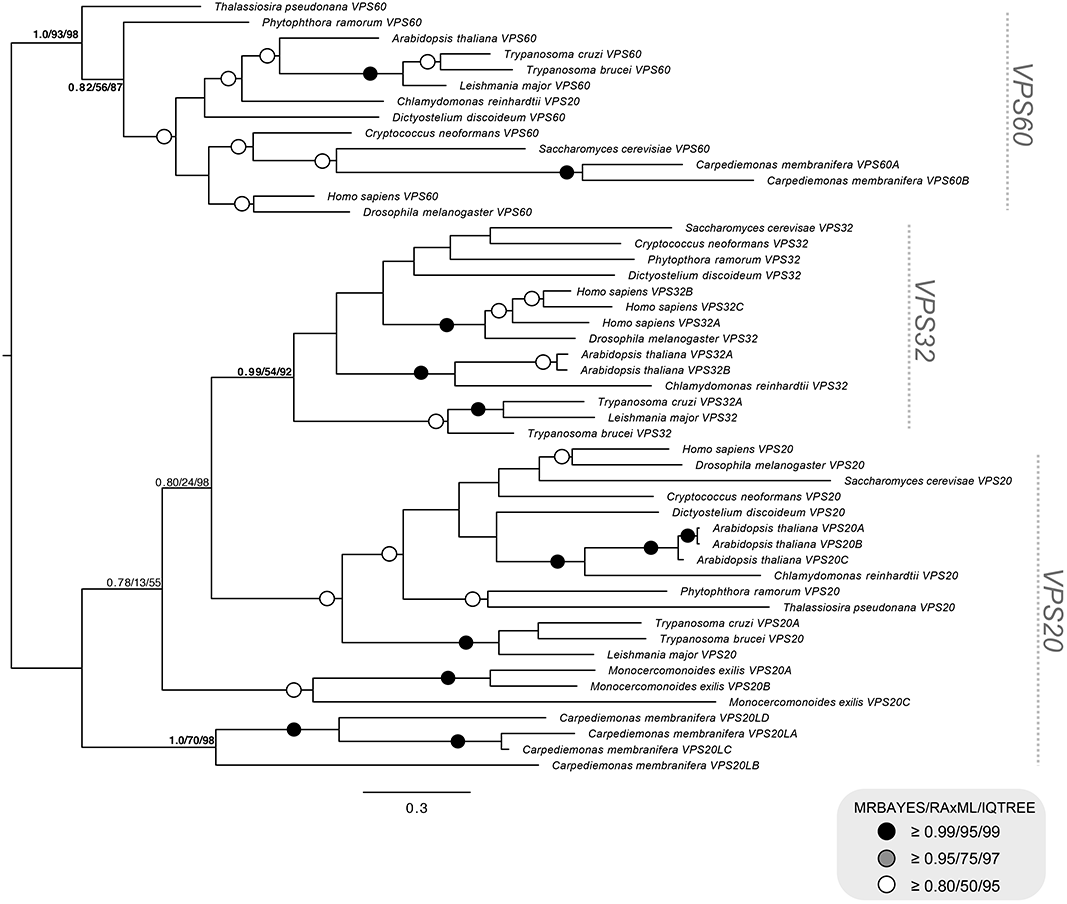
Phylogenetic analyses of the individual VPS20-SNF7 family proteins from ESCRTIII and ESCRTIIIA sub-complexes which depicts pan-eukaryotic VPS20/32/60 with *Carpediemonas membranifera* SNF7 family proteins used as landmark representative for Fornicata. Tree inference was carried out using both BI and ML analyses. RAxML best model was determined to be LG+G+F while IQ-TREE ModelFinder determined an equivalent LG+G4+F. Two of the identified SNF7 sequences from *Carpediemonas membranifera* clustered clearly with VPS60 whereas the remainder neither grouped with VPS20 or VPS32 and therefore were determined to be VPS20L proteins in all tree topologies

**Figure S3.**
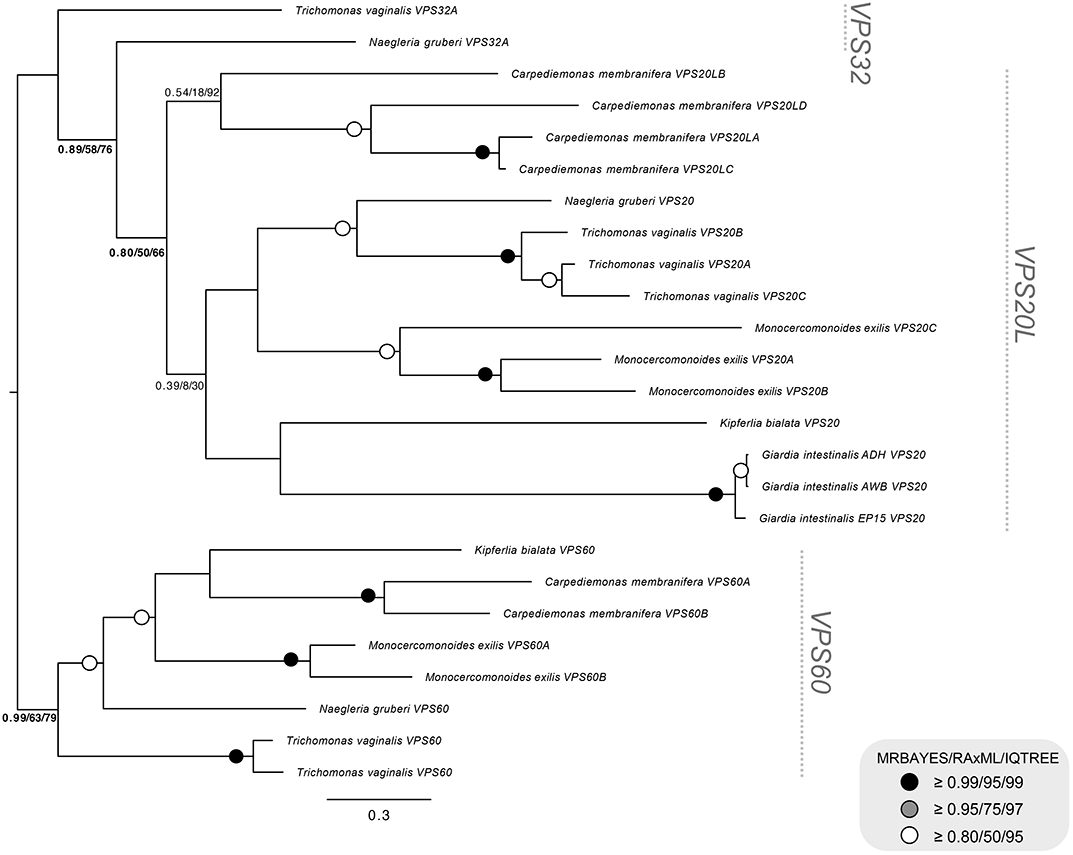
Phylogenetic analyses of the individual VPS20-SNF7 family proteins from ESCRTIII and ESCRTIIIA sub-complexes depicts a Fornicata specific tree with well characterized Excavata representatives *Monocercomonoides exilis, Trichomonas vaginalis,* and *Naegleria gruberi* where no identified diplomonad SNF7 sequences grouped with VPS60. All identified *Giardia* SNF7 sequences grouped with VPS20 from other metamonads and therefore were also determined to be VPS20L sequences. Both trees were rooted at ESCRTIII-VPS60 highlighted in red (*Leung* et al., 2008).

**Figure S4.**
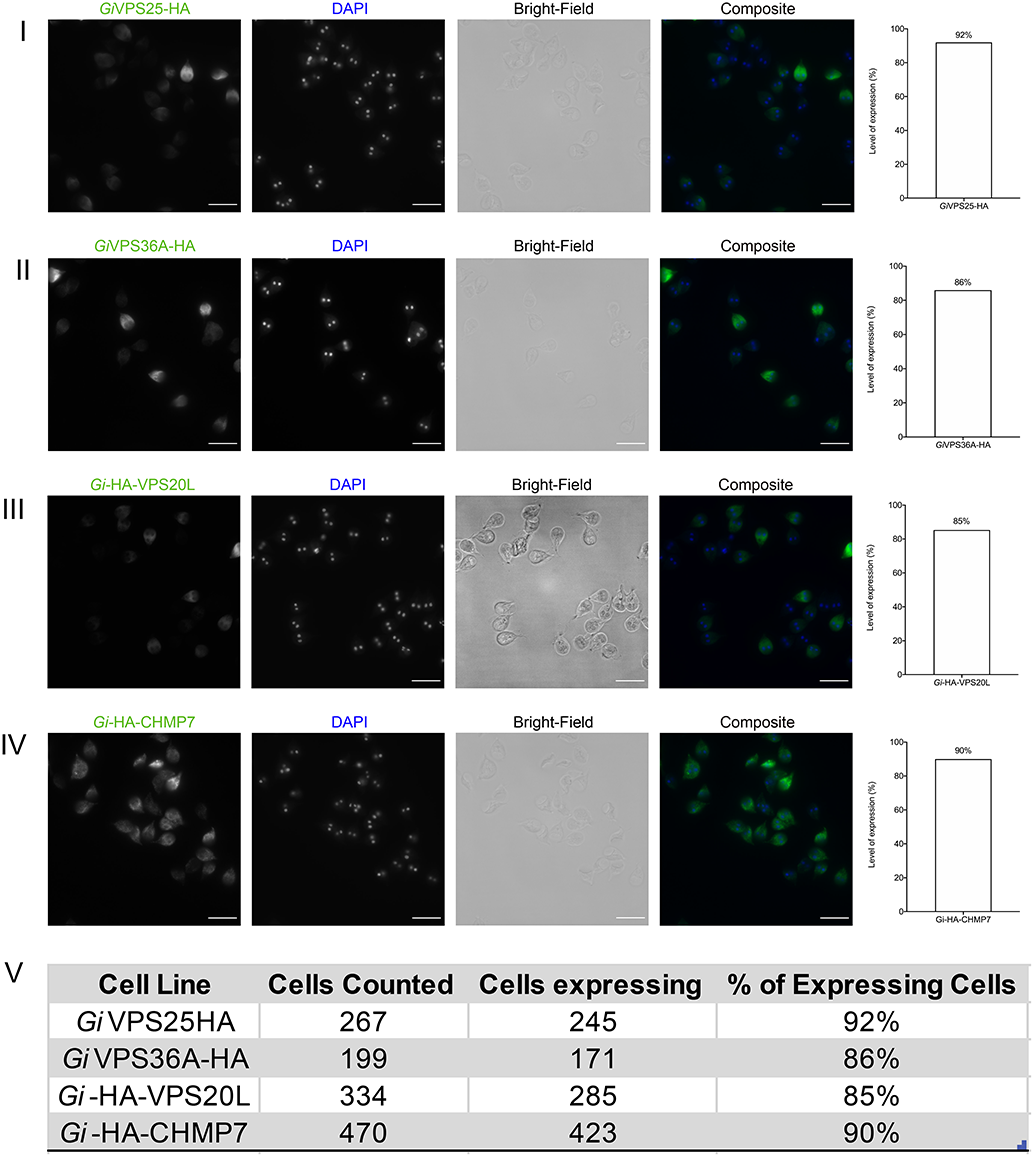
Population-level expression analysis of epitope-tagged ESCRT subunits. (I) *Gi*VPS25HA is expressed in 92% of screened cells. (II) *Gi*VPS36A-HA is expressed in 86% of screened cells. (III) *Gi-*HA-VPS20L is expressed in 85% of cells while (IV) *Gi-*HA-CHMP7 is expressed in 90% of the cells. (V) Detailed results used for quantification. All scale bars: 20 μm.

**Figure S5.**
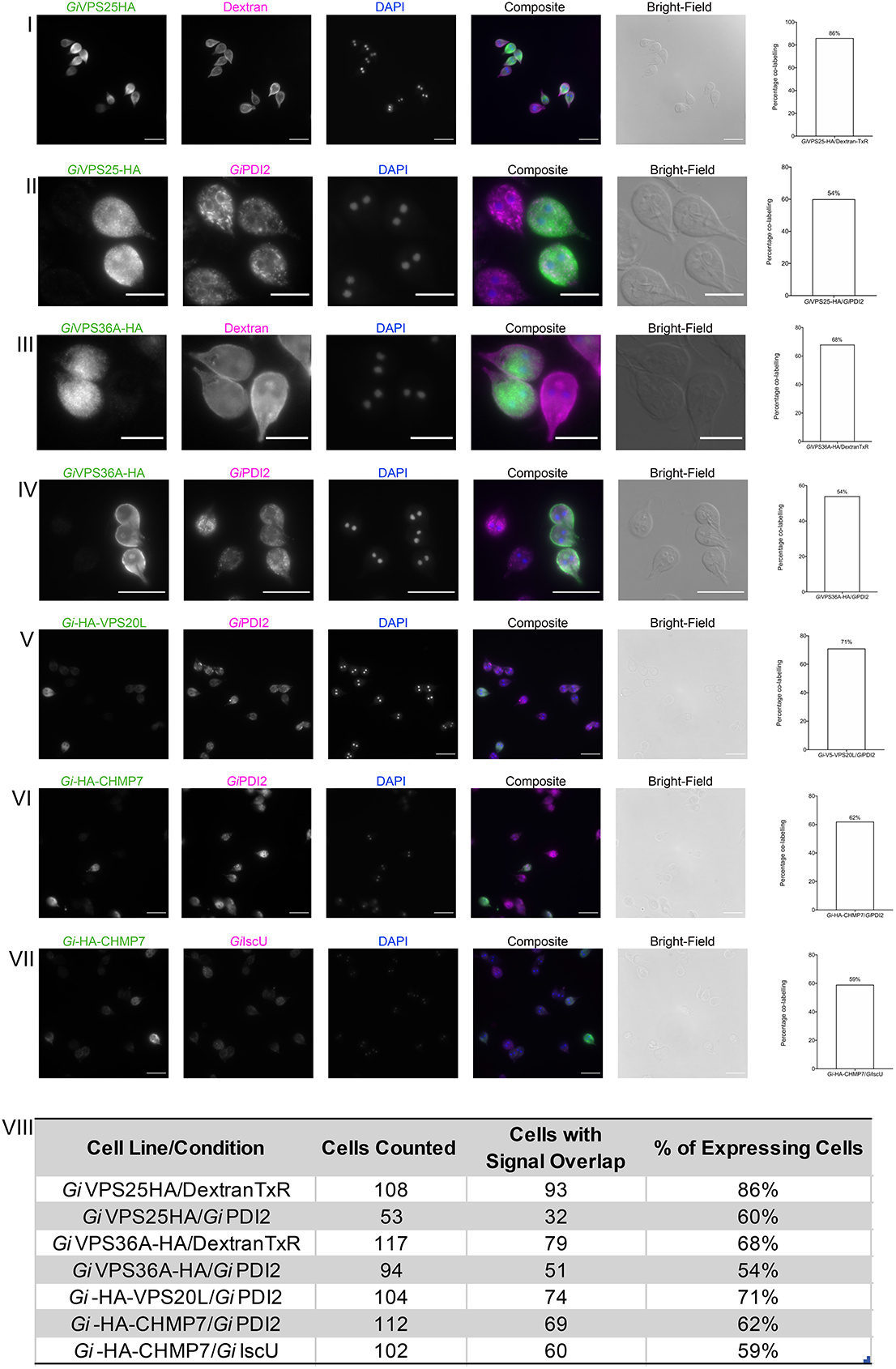
Population-level analysis of cells co-labelled for ESCRT subunits and selected subcellular markers. (I) *Gi*VPS25-HA with Dextran TexasRed and (II) with *Gi*PDI2. (III) *Gi*VPS36A-HA with Dextran TexasRed and (IV) with *Gi*PDI2. (V) *Gi-*HA-VPS20L with *Gi*PDI2. (VI) *Gi-*HA-CHMP7 with *Gi*PDI2 and (VII) when counterstained for *Gi*IscU. (VIII) Detailed results used for signal overlap quantification. Scale bars: (I, V-VII) 20 μm and (II-IV) 10 μm.

**Figure S6.**
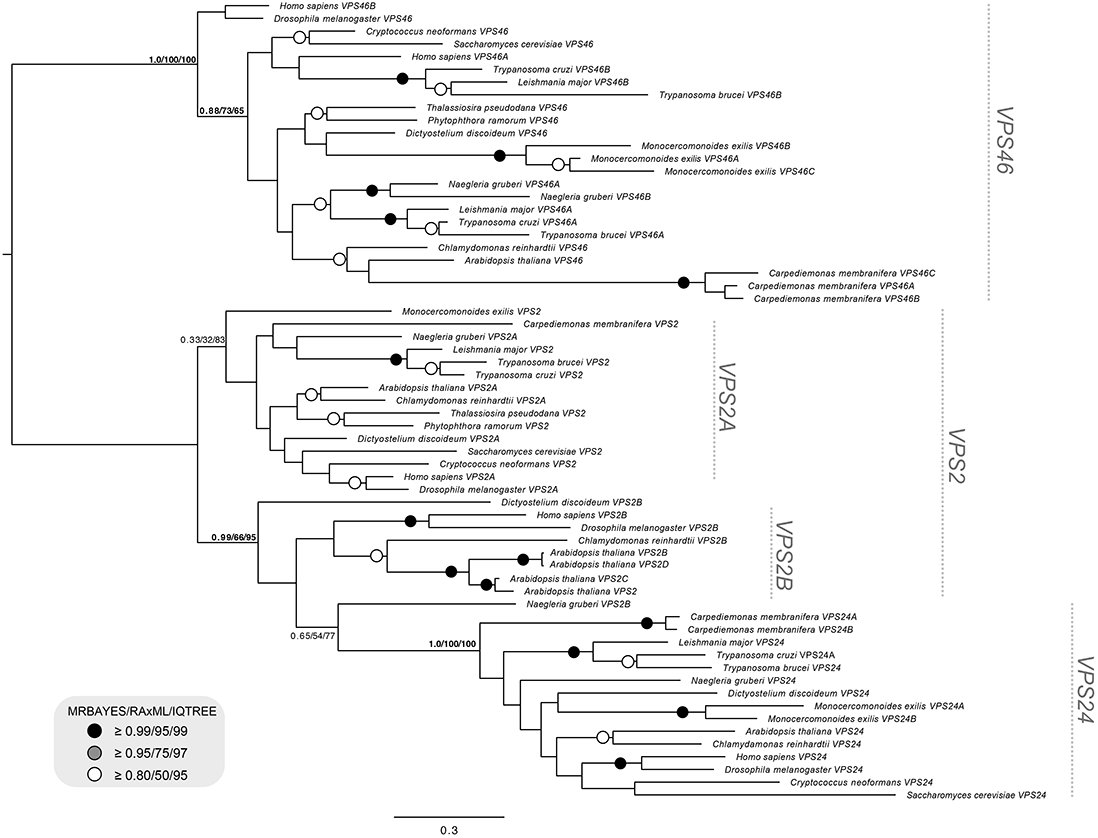
Phylogenetic analyses of the individual SNF7-VPS2 family proteins from ESCRTIII and ESCRTIII-A sub-complexes depicts pan-eukaryotic VPS2/24/46 with *Carpediemonas membranifera* VPS2 family proteins used as landmark representative for Fornicata. Tree inference was carried out using both Bayesian Inference and Maximum Likelihood analyses. RAxML best model was determined to be LG+G+F while IQ-TREE ModelFinder determined an equivalent LG+G4+F. *Carpediemonas membranifera* was determined to have all three components with strong backbone support for VPS24 (1.0/100/100) and VPS46 (1.0/100/100).

**Figure S7.**
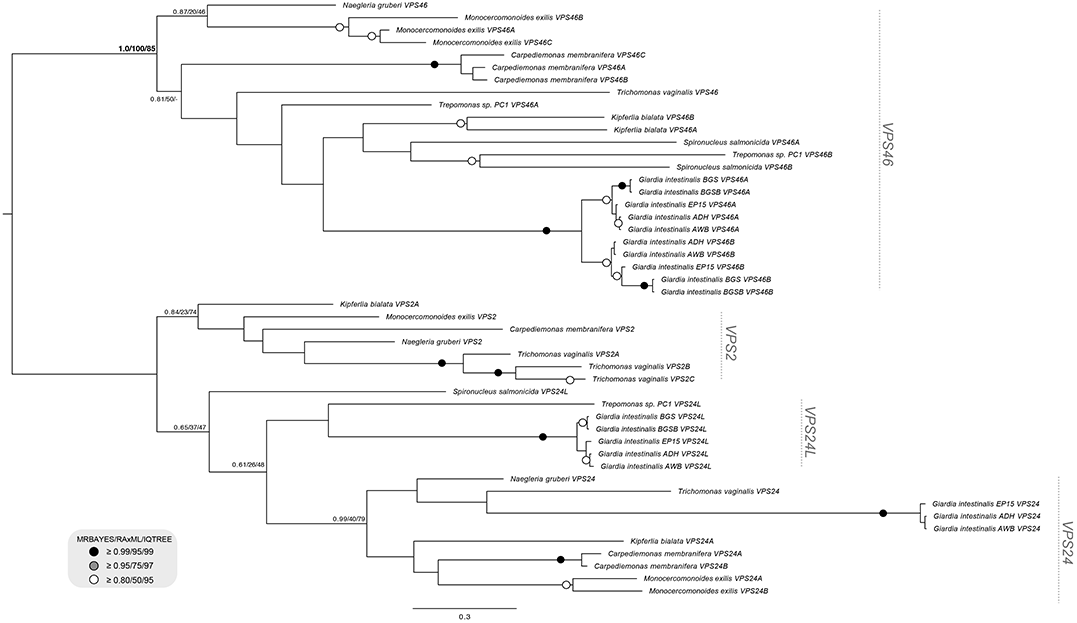
Phylogenetic analyses of the individual SNF7-VPS2 family proteins from ESCRTIII and ESCRTIII-A sub-complexes depicts a Fornicata specific tree with well characterized Excavata representatives *Monocercomonoides exilis., Trichomonas vaginalis,* and *Naegleria gruberi* in order to classify divergent diplomonad sequences. VPS2 family proteins identified in the diplomonads grouped with both VPS24 and VPS46 with duplication event pointing in *Giardia* spp. VPS46 yielding two paralogues, VPS46A and VPS46B. An additional set of VPS2 family proteins which neither grouped clearly with VPS2 or VPS24 and therefore were determined to be VPS24 like proteins. Tree was rooted at ESCRTIII-VPS46 clade (Leung et. al. 2008).

**Figure S8.**
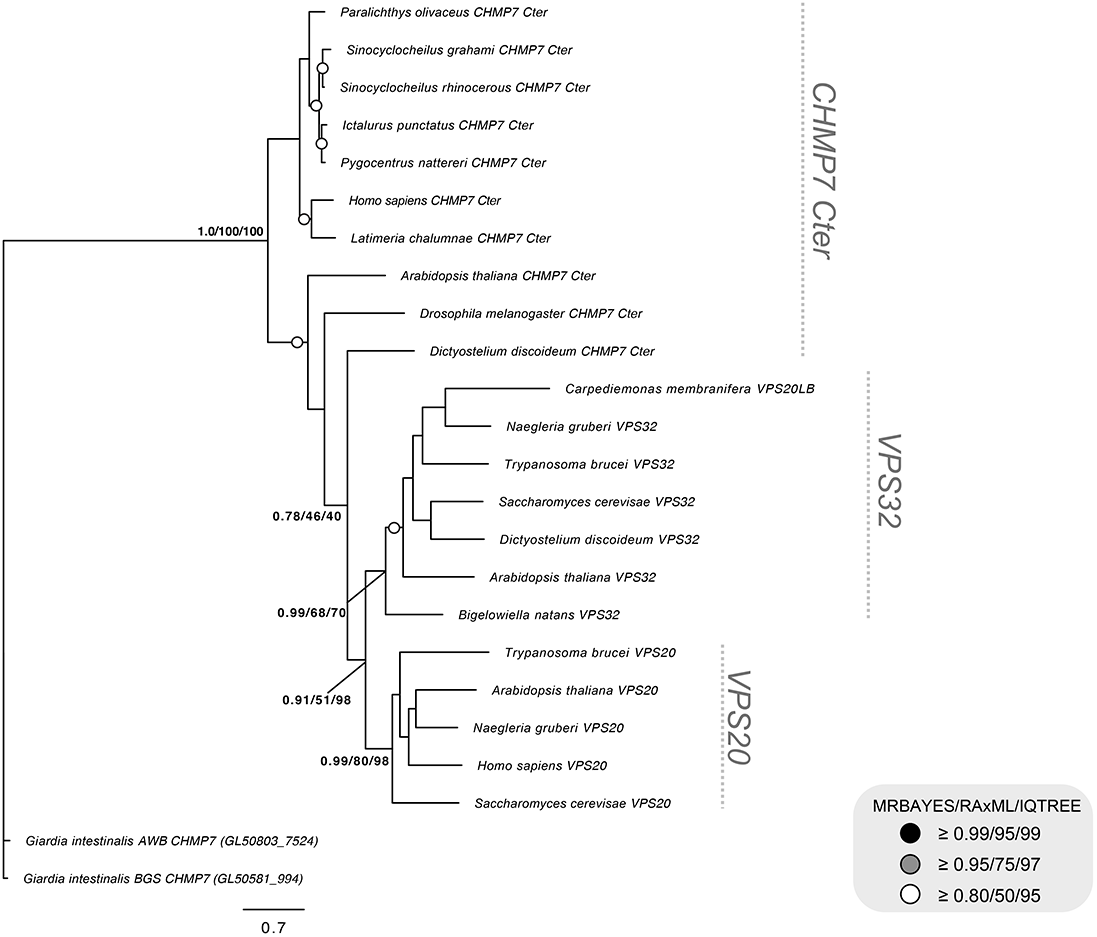
Phylogenetic analyses of the *Gi*AWBCHMP7 and *Gi*BGSCHMP7 against SNF7 components and CHMP7 c-terminus. Unrooted phylogenetic analyses of the identified *Giardia* CHMP7 against CHMP7 c-termini from various pan-eukaryotic lineages and previously its previously proposed homology to VPS20/32 SNF7 show exclusion of the *Giardia* proteins with MRBAYES, RAxML, and IQTREE support 1.0/100/100

**Figure S9.**
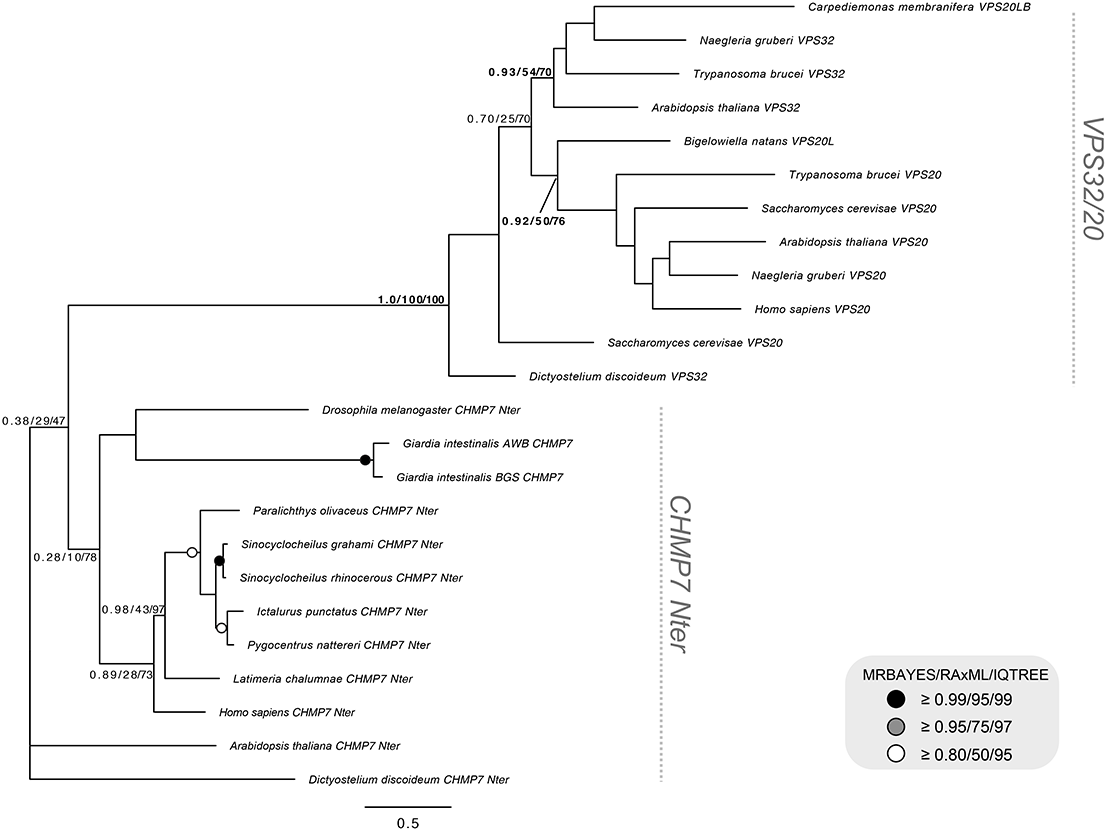
Phylogenetic analyses of the *Gi*AWBCHMP7 and *Gi*BGSCHMP7 against SNF7 components and CHMP7 N-terminus. Unrooted phylogenetic analyses of the identified CHMP7 against CHMP7 n-terminus and VPS20/32 SNF7 show inclusion of the *Giardia* proteins within the pan-eukaryotic CHMP7 N-termini clade away from the VPS20/32 clade with the support of 1.0/100/100

**Figure S10.**
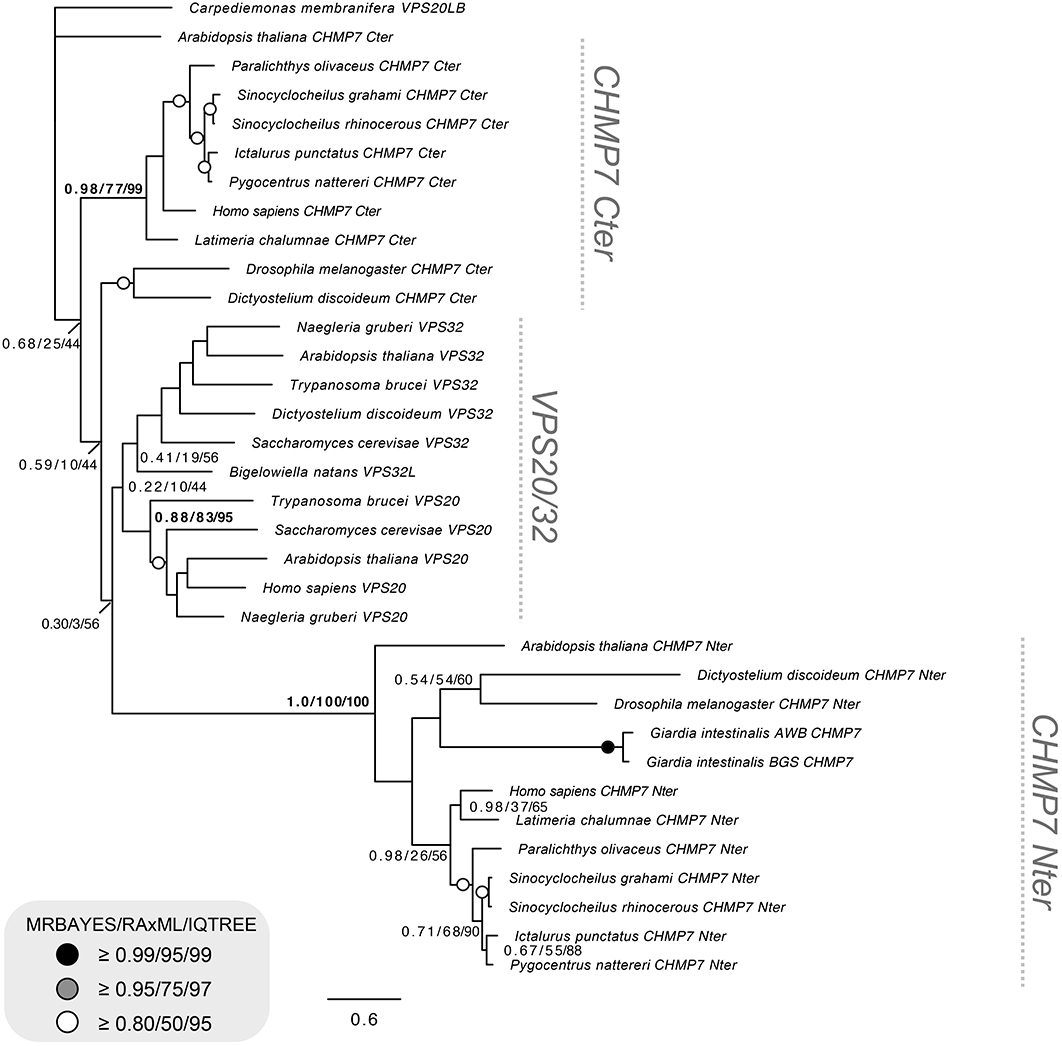
Phylogenetic analyses of the pan-eukaryotic CHMP7 N-termini against SNF7 VPS20/32 and CHMP7 C-terminus. Unrooted phylogenetic analyses of pan-eukaryotic CHMP7 N-termini form a clade with exclusion to pan-eukaryotic CHMP7 C-termini and pan-eukaryotic VPS20 and VPS32 contrary to the previously proposed homology to SNF7 with MRBAYES, RAxML, and IQTREE support of 1.0/100/100.

**Figure S11.**
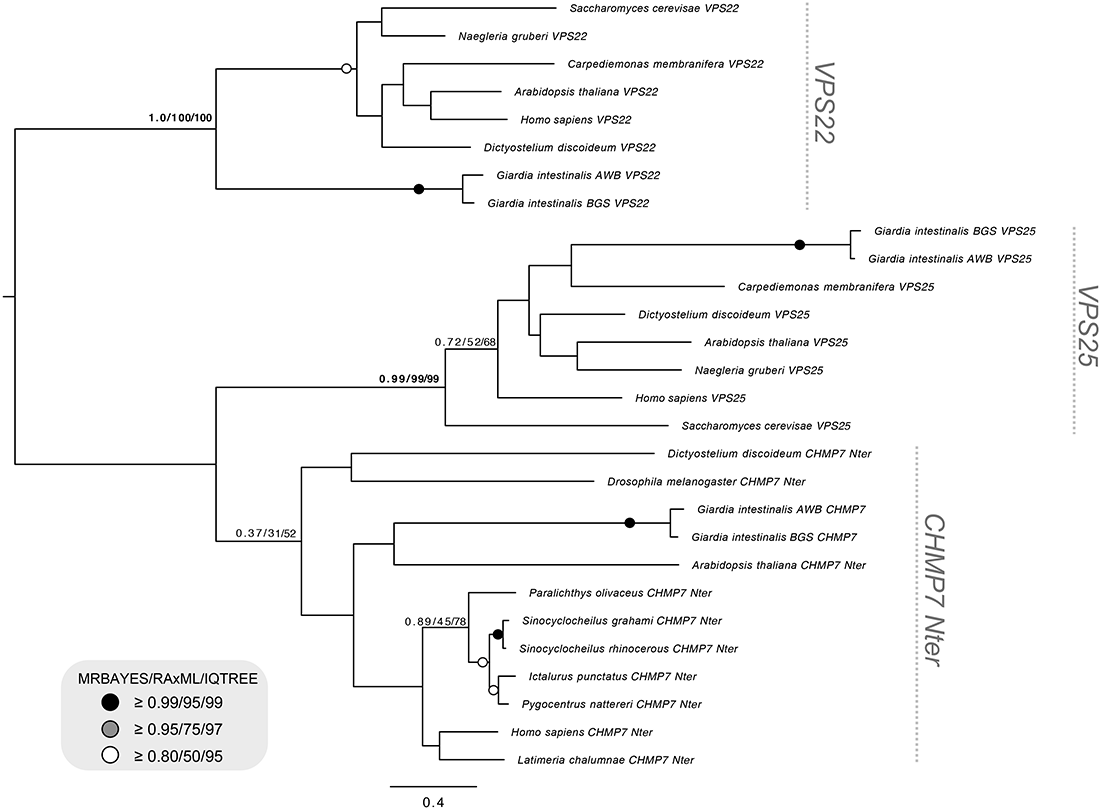
Phylogenetic analyses of the *Gi*AWBCHMP7 and *Gi*BGSCHMP7 and pan-eukaryotic and pan-eukaryotic CHMP7 N-termini against pan-eukaryotic VPS25 orthologs. HHPRED analyses (See Supplementary Table S3) of the *Giardia* CHMP7 proteins showed closest homology to ESCRTII-VPS25 and therefore were phylogenetically tested to ensure that these were in fact not additional paralogs of *Giardia* VPS25. Identified CHMP7 proteins were in fact not paralogs of the *Giardia* VPS25 which grouped in the separate VPS25 clade with the backbone support of 0.99/99/99 to the exclusion of CHMP7 clade. Tree was rooted onto ESCRTII-VPS22 pan-eukaryotic proteins.

### Videos

**Supplementary Video 1.** Video reconstruction of *Gi*VPS25-HA confocal imaging analyses and 3-dimensional rendering of subcellular localization, as depicted in Figure 3A.

**Supplementary Video 2.** Video reconstruction of *Gi*VPS25-HA confocal imaging analyses and 3-dimensional rendering of intracellular co-localization with Dextran-TexasRed, as depicted in Figure 3B.

**Supplementary Video 3.** Video reconstruction of *Gi*VPS25-HA confocal imaging analyses and 3-dimensional rendering of intracellular co-localization with *Gi*PDI2, as depicted in Figure 5B.

**Supplementary Video 4.** Video reconstruction of *Gi*VPS36-HA confocal imaging analyses and 3-dimensional rendering of intracellular localization, as depicted in Figure 4A.

**Supplementary Video 5.** Video reconstruction of *Gi*VPS36-HA confocal imaging analyses and 3-dimensional rendering of intracellular co-localization with Dextran-TexasRed, as depicted in Figure 4B.

**Supplementary Video 6.** Video reconstruction of *Gi*VPS36-HA confocal imaging analyses and 3-dimensional rendering of intracellular co-localization with *Gi*PDI2, as depicted in Figure 5C.

**Supplementary Video 7.** Video reconstruction of *Gi*HA-VPS20L confocal imaging analyses and 3-dimensional rendering of intracellular localization, as depicted in Figure 5A.

**Supplementary Video 8.** Video reconstruction of *Gi*HA-VPS20L confocal imaging analyses and 3-dimensional rendering of intracellular co-localization with *Gi*PDI2, as depicted in Figure 5D.

**Supplementary Video 9.** Video reconstruction of *Gi*HA-CHMP7 confocal imaging analyses and 3-dimensional rendering of intracellular localization, as depicted in Figure 8A

**Supplementary Video 10.** Video reconstruction of *Gi*HA-CHMP7 confocal imaging analyses and 3-dimensional rendering of intracellular co-localization with *Gi*PDI2, as depicted in Figure 8B

**Supplementary Video 11.** Video reconstruction of *Gi*HA-CHMP7 confocal imaging analyses and 3-dimensional rendering of intracellular co-localization with *Gi*IscU, as depicted in Figure 8C

### Tables

**Table S1.** Pan eukaryotic ESCRT queries and databases used for retrieval of up-to-date genomes and transcriptomes for homology searching and phylogenetic analyses

**Table S2.** Identified ESCRT components in Fornicata genomes and transcriptomes identified and validated using HMMER, tBLASTn, BLASTP, HHPRED, and CDD domain searches.

**Table S3.** HHPred analyses of pan-eukaryotic, including *Giardia*, CHMP7 N-termini.

**Table S4.** Oligonucleotides used for the cloning of *Giardia* ESCRT subunits of interest.

